# Validation studies and multi-omics analysis of Zhx2 as a candidate quantitative trait gene underlying brain oxycodone metabolite (oxymorphone) levels and behavior

**DOI:** 10.1101/2024.08.30.610534

**Authors:** William B. Lynch, Sophia A. Miracle, Stanley I. Goldstein, Jacob A. Beierle, Rhea Bhandari, Ethan T. Gerhardt, Ava Farnan, Binh-Minh Nguyen, Kelly K. Wingfield, Ida Kazerani, Gabriel A. Saavedra, Olga Averin, Britahny M. Baskin, Martin T. Ferris, Christopher A. Reilly, Andrew Emili, Camron D. Bryant

**Author notes:** Corresponding Author: Camron D. Bryant, Ph.D., Laboratory of Addiction Genetics Department of Pharmaceutical Sciences Center for Drug Discovery, Northeastern University 140 The Fenway, X138, Boston, MA USA 02115, E.

## Abstract

Sensitivity to the subjective reinforcing properties of opioids has a genetic component and can predict addiction liability of opioid compounds. We previously identified *Zhx2* as a candidate gene underlying increased brain concentration of the oxycodone (**OXY**) metabolite oxymorphone (**OMOR**) in BALB/cJ (**J**) versus BALB/cByJ (**By**) females that could increase OXY state-dependent reward. A large structural intronic variant is associated with a robust reduction of Zhx2 expression in J mice, which we hypothesized enhances OMOR levels and OXY addiction-like behaviors. We tested this hypothesis by restoring the *Zhx2* loss-of-function in Js (**MVKO**) and modeling the loss-of-function variant through knocking out the *Zhx2* coding exon (**E3KO**) in Bys and assessing brain OXY metabolite levels and behavior. Consistent with our hypothesis, Zhx2 E3KO females showed an increase in brain OMOR levels and OXY-induced locomotor activity. However, contrary to our hypothesis, state-dependent expression of OXY-CPP was decreased in E3KO females and increased in E3KO males. We also overexpressed Zhx2 in the livers and brains of Js and observed Zhx2 overexpression in select brain regions that was associated with reduced OXY state-dependent learning. Integrative transcriptomic and proteomic analysis of E3KO mice identified astrocyte function, cell adhesion, extracellular matrix properties, and endothelial cell functions as pathways influencing brain OXY metabolite concentration and behavior. These results support *Zhx2* as a quantitative trait gene underlying brain OMOR concentration that is associated with changes in OXY behavior and implicate potential quantitative trait mechanisms that together inform our overall understanding of *Zhx2* in brain function.

## INTRODUCTION

Opioid Use Disorder **(OUD)** remains a major public health epidemic in the United States. While widespread misuse of fentanyl is driving the current crisis in opioid overdose deaths[1–3], prescription opioid misuse from compounds like Oxycodone (**OXY**), one of the most widely prescribed and misused prescription opioids, continues to fuel the opioid epidemic[4–7]. Despite existing medications in treating OUD, there is an urgent need for novel treatment and prevention strategies[8,9] and further studies assessing sex differences in opioid response[10,11]. OUD heritability is estimated at 50%[12–14]. However, the genetic and molecular mechanisms underlying OUD risk remains poorly understood[15]. A more complete understanding of the genetic basis of OUD-associated molecular and behavioral traits could inform molecular mechanisms and thus more effective and tailored treatments.

Phenotypic variation in opioid pharmacokinetic measurements has a strong genetic component that can dramatically influence physiological and subjective responses to opioids. The most well-known example is Cytochrome P450 2D6 (**CYP2D6**) which converts the inactive prodrug codeine into its active metabolite morphine[16]. CYP2D6 also similarly converts OXY to a highly active metabolite oxymorphone (**OMOR**) alongside the inactive metabolite noroxycodone (**NOR**). Individual enzymatic activity differences can influence clinical OXY analgesia[17–19]. OMOR has a 40-fold increase in mu-opioid receptor affinity versus OXY and therefore could enhance acute and conditioned OXY responses[20–22]. Additionally, sex differences in OXY to OMOR metabolism are reported and could underlie sex-specific OXY responses[23,24]. However, whether OMOR metabolite levels contribute to OXY behaviors remains largely unexplored, although some studies indicate OMOR metabolite effects on OXY analgesic and addictive properties[19,25].

Rodent quantitative genetics offers the ability to discover and validate candidate variants within the same species[26,27]. Reduced complexity crosses between near-isogenic substrains can expedite identifying causal genes and variants underlying complex trait variation[28,29]. BALB/cJ **(J)** and BALB/cByJ **(By)** have a drastically reduced genetic complexity versus classical inbred strains, yet differ in multiple behavioral and molecular phenotypes[29–31]. We found that J females showed increased OXY state-dependent reward learning and concomitantly increased brain OMOR levels compared to By females[32]. Quantitative trait locus mapping of OXY traits and gene expression (**eQTL**) in a BALB/c reduced complexity cross identified Zinc-Finger and Homeoboxes 2 (**Zhx2**) as the top candidate gene regulating brain OMOR concentration[32]. The likely variant mediating the eQTL for Zhx2 expression is a mouse endogenous retroviral element (**MERV**) insertion specific to Js[33,34]. However, whether or not Zhx2 expression and function are linked to brain OMOR levels and OXY behavior remains a key question.

*Zhx2* codes for a transcription factor acting largely as a transcriptional repressor. *Zhx2* is historically associated with cancer and influences lipid metabolism, inflammatory processes, and cellular differentiation[35,36]. Notably, *Zhx2* function is associated with altered liver CYP enzyme expression in a sex-dependent manner[37]. Rodent Cyp2d orthologs are active in both the liver and brain and influence OXY antinociception[17,38]. Furthermore, we previously observed increased Cyp2d11 transcript levels in J vs. By mouse brains[32]. Therefore, we hypothesized that decreased Zhx2 in Js could upregulate Cyp2d brain and/or liver expression to increase OMOR levels and OXY addiction-like behaviors.

The primary goal of this study was to validate *Zhx2* as a quantitative trait gene underlying the female-specific increase in brain OMOR levels[32] and associate these changes with OXY behavior using reciprocal gene editing on each of the two BALB/c substrain backgrounds. We hypothesized that decreasing Zhx2 expression would increase CYP2D expression, increase brain OMOR concentration, and increase OXY addiction-like behaviors in females and that the reciprocal results would be observed when increasing Zhx2.

## MATERIALS AND METHODS

### Mice

CRISPR/Cas9 gene editing on each BALB/c genetic background was performed by The Jackson Laboratory (**JAX,** Bar Harbor, ME). Mice were shipped and bred in-house at Boston University. Details on transgenic mouse generation and breeding are provided in **Supplementary Materials and Methods.** All experiments were conducted in strict accordance with National Institute of Health guidelines for the Care and Use of Laboratory Animals and approved by the Boston University Institutional Animal Care and Use Committee.

### Adenoassociated virus (AAV) generation and administration

All AAVs were custom constructed and produced by Vector Biolabs (Malburn, PA), which included overexpression **(OE)** and control **(CTR)** constructs for the brain and liver respectively. Each mouse was injected at least 3 weeks prior to behavioral experiments. Details on viral administration are in **Supplementary Materials and Methods.**

### Conditioned place preference (CPP)

CPP was performed as described[39]. Mice received 1.25 mg/kg OXY intraperitoneal **(i.p.)** on respective days. A description of the CPP procedure can be found in **Supplementary Materials and Methods.**

### Tissue Extractions/Perfusions

Brains and livers were harvested post-CPP. Left liver lobes and brain hemispheres were flash frozen in 2-methylbutane and stored at -80degC while right lobes and hemispheres were stored in RNA-later for 2-4 days before -80degC storage[40]. A subset of brains were transcardially perfused with 1X phosphate buffered saline **(PBS)** and 4% paraformaldehyde **(PFA)** for immunohistochemistry.

### Real Time quantitative PCR (qPCR) of RNA expression

RNA from isolated tissues was extracted via Trizol homogenization and chloroform precipitation. Samples were treated with DNase (Qiagen, Hilden, Germany) and eluted using a RNeasy kit (Qiagen). RNA was measured and diluted to 100 ng/ul for cDNA library generation using random primers (ThermoFisher, Waltham, MA). cDNA was diluted 1:10 and RNA was quantified via a QuantStudio 12K Flex Real-Time PCR System (ThermoFisher). Target RNA C_T_ values were normalized to within sample beta-actin C_T_ values. Data was transformed and compared between conditions using the 2^-ΔΔC^ method[41].

### Western blotting for ZHX2 protein

Proteins were extracted and quantified via a BCA assay. 30 ug of protein in each sample was run on a Precast Midi Protein Gel (Bio-Rad, Hercules, CA) and transferred to a membrane blot overnight at 4degC. Blots were blocked for 1 hr in 5% condensed milk dissolved in Tris-Buffered Saline + 0.1% Tween-20 **(TBST).** Blots were incubated in a ZHX2 primary antibody overnight (GeneTex; 1:4000) at 4degC and then a secondary antibody (Jackson Immunoresearch; 1:10K) for 1 hr. Blots were visualized with a Clarity Western ECL Substrate (Bio-Rad) and band intensities were quantified using ImageJ[42]. Intensity percentages of each sample were calculated and normalized to the intensity percentages generated from the Ponceau S stain with each sample before factor comparison[43].

### Immunohistochemistry of brain Zhx2 OE spread

To assess viral spread of AAVs for brain Zhx2 OE, 30 um coronal slices from brains of virally injected mice were stained with a primary ZHX2 antibody (GeneTex; 1:750) and imaged using a Confocal Microscope. A detailed description is provided in **Supplementary Materials and Methods.**

### OXY metabolite analysis

Left brain hemispheres were weighed and homogenized in molecular grade water. Homogenate was shipped on dry ice to the University of Utah Center for Human Toxicology and OXY, NOR, and OMOR brain concentrations were quantified as previously described[32,44]. Concentrations were normalized to brain weight and compared between conditions.

### RNA-sequencing

RNA from right brain hemispheres was extracted and cDNA libraries were prepared and sequenced by the University of Chicago Genomics Facility using an Illumina NovaSeqX (Illumina, San Diego, CA) with a sequencing depth of 50-60 million paired end reads per sample as described previously[45]. All read processing and comparison was performed using GALAXY[46]. Reads were trimmed for quality using Trimmomatic[47] and aligned to the mm10 reference genome using STAR[48]. Reads were counted against the GRCm38 refence genome using FeatureCounts[49] and compared between conditions using edgeR[50]. Genes without at least 10 reads per million in at least 3 samples were excluded from analysis.

### Proteomic mass-spectrometry

All sample preparation and mass-spectrometry was performed as previously described[51], with additional details provided in **Supplementary Materials and Methods.** Subsequent data filtering, normalization, and analysis were performed using the Omics Notebook pipeline[52].

### Multi-Omics pathway enrichment and visualization

Pathway enrichment analysis was performed using the R package fgsea[53]. Pathway enrichment visualization and leading-edge gene interactions were performed using the R package enrichplot[54] along with the STRING-db database[55]. A detailed description of these methods is in **Supplementary Materials and Methods.**

### Statistics

All statistics were conducted using R: *A language and environment for Statistical Computing.* Detailed descriptions are in **Supplementary Materials and Methods.**

## RESULTS

### Effect of *Zhx2* MERV gene deletion on OXY metabolite levels and behaviors in female and male J mice

A list of antibodies for western blotting and IHC is provided in **Table S1** while a list of qPCR primers is provided in **Table S2**. Sample sizes for main figures are provided in **Figure S1** while sample sizes for supplemental figures are present in **Figure S2**.

In order to test *Zhx2* MERV necessity for increasing brain OMOR levels and addiction-like behaviors, we used CRISPR gene editing to remove the MERV insertion from Js (**Fig. S3, Fig. S4A**). Subsequent protein validation indicated MERV knockout (**MVKO**) restored brain ZHX2 expression to a By-like level (**Fig. S3B**). However, ZHX2 expression restoration did not significantly affect brain OXY, NOR, or OMOR levels in both sexes (**Fig. S3C-H**). Complimenting the metabolite results, there were no significant genotype differences in distance or time spent phenotypes in CPP in MVKOs (**Fig. S5**). Thus, MVKO was not sufficient to reduce brain OMOR levels or OXY behaviors in contrast to our original hypothesis.

### Effect of Zhx2 liver overexpression on liver Cyp2d expression, OXY brain metabolite levels, and OXY behaviors in J mice

While the CRISPR-mediated ZHX2 increase in Js via MVKO was insufficient to alter OXY metabolite levels and behavior, we overexpressed Zhx2 in select tissues in the original J parental strain as another means to rescue the *Zhx2* MERV loss of function and determine its effect on OXY metabolite levels and behavior. Here, we selectively overexpressed Zhx2 in the liver vs. brain because of both tissues’ importance in OXY metabolism and behavior. We increased liver Zhx2 expression without modifying brain Zhx2 expression in J mice (**Fig. 1A-B**; plasmid map in **Fig. S6A**). We then analyzed expression of a subset of 3 out of 9 total Cyp2d mouse genes, selected based on suspected relevance to *Zhx2* and opioid metabolism[32,37,56], in the liver between viral conditions. Zhx2 liver OE increased liver Cyp2d22 mRNA expression in J females (**Fig. 1C**) whereas it decreased expression in J males (**Fig. 1E**). There was no change in Cyp2d11 or Cyp2d10 expression (**Fig. S7**). Despite liver Cyp changes, no changes in brain OMOR concentration were observed (**Fig. 1D, F**). Interestingly, Zhx2 OE significantly decreased brain OXY concentration exclusively in males (**Fig. S8**). Aside from some non-selective locomotor differences, Zhx2 liver OE had no obvious impact on OXY behavior (**Fig. S9**). A detailed description of these results is provided in **Supplementary Results.** Overall, while liver Zhx2 overexpression affected hepatic Cyp2d22 transcript levels, its effects on brain OXY metabolite levels and behaviors were limited and non-selective.

**Figure 1.**
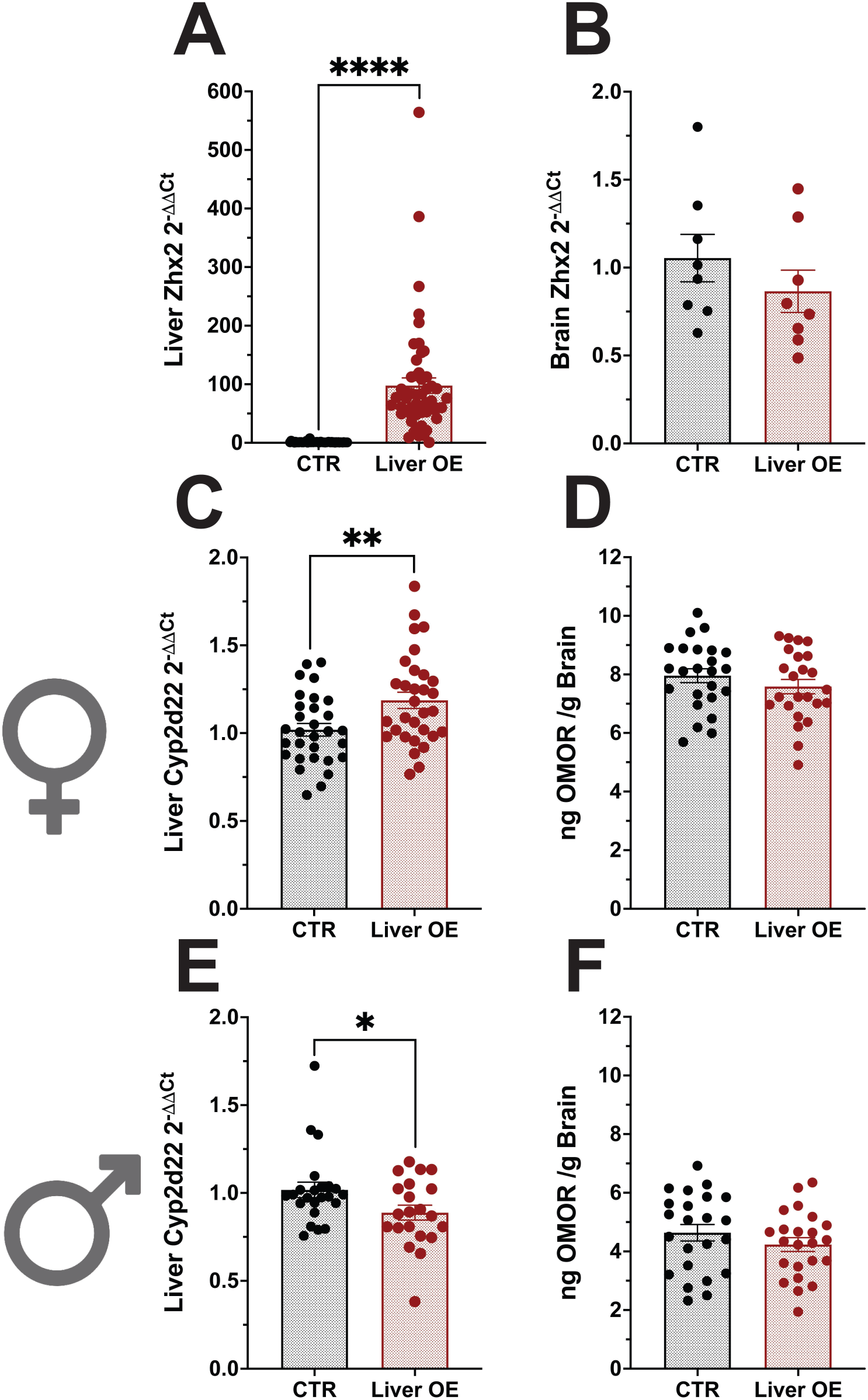
Quantification of Zhx2 Liver OE, Cytochrome P450 transcripts, and brain OMOR in BALB/cJ (J) mice following OXY-CPP (1.25 mg/kg, i.p.). **(A):** Liver Zhx2 RNA expression following liver OE versus controls (t_51.013_ = 7.28, ****p < 0.0001). **(B):** Brain Zhx2 RNA expression following liver OE in J females and males (t_14_ = 1.05, p = 0.314). **(C):** Liver Cyp2d22 RNA expression following liver OE in J females (t_59_ = 2.84, **p < 0.01). **(D):** Brain OMOR concentration following liver OE in J females (t_46_ = 1.10, p = 0.276). **(E):** Liver Cyp2d22 RNA expression following liver Zhx2 OE in J males (t_42_ = 2.12, *p < 0.05). (F): Brain OMOR brain concentrations following liver OE in J males (t_44_ = 1.11, p = 0.275).

### Effect of Zhx2 brain overexpression on OXY behaviors in J mice

We next sought to overexpress Zhx2 in J mouse brains via AAV intracerebroventricular (**i.c.v.**) administration (plasmid map in **Fig. S6B**). Immunohistochemical analysis indicated marked, localized increases in fluorescent staining specifically within the lateral septum (**LS**), bed nucleus of the stria terminalis (**BNST**), and hippocampal CA3 subregion of OE mice compared to controls (**Fig 2A**, image sites in **Fig. S10**). We saw no changes in OXY metabolite concentrations (data not shown), which in hindsight would be expected given the highly localized as opposed to widespread pattern of Zhx2 OE.

**Figure 2.**
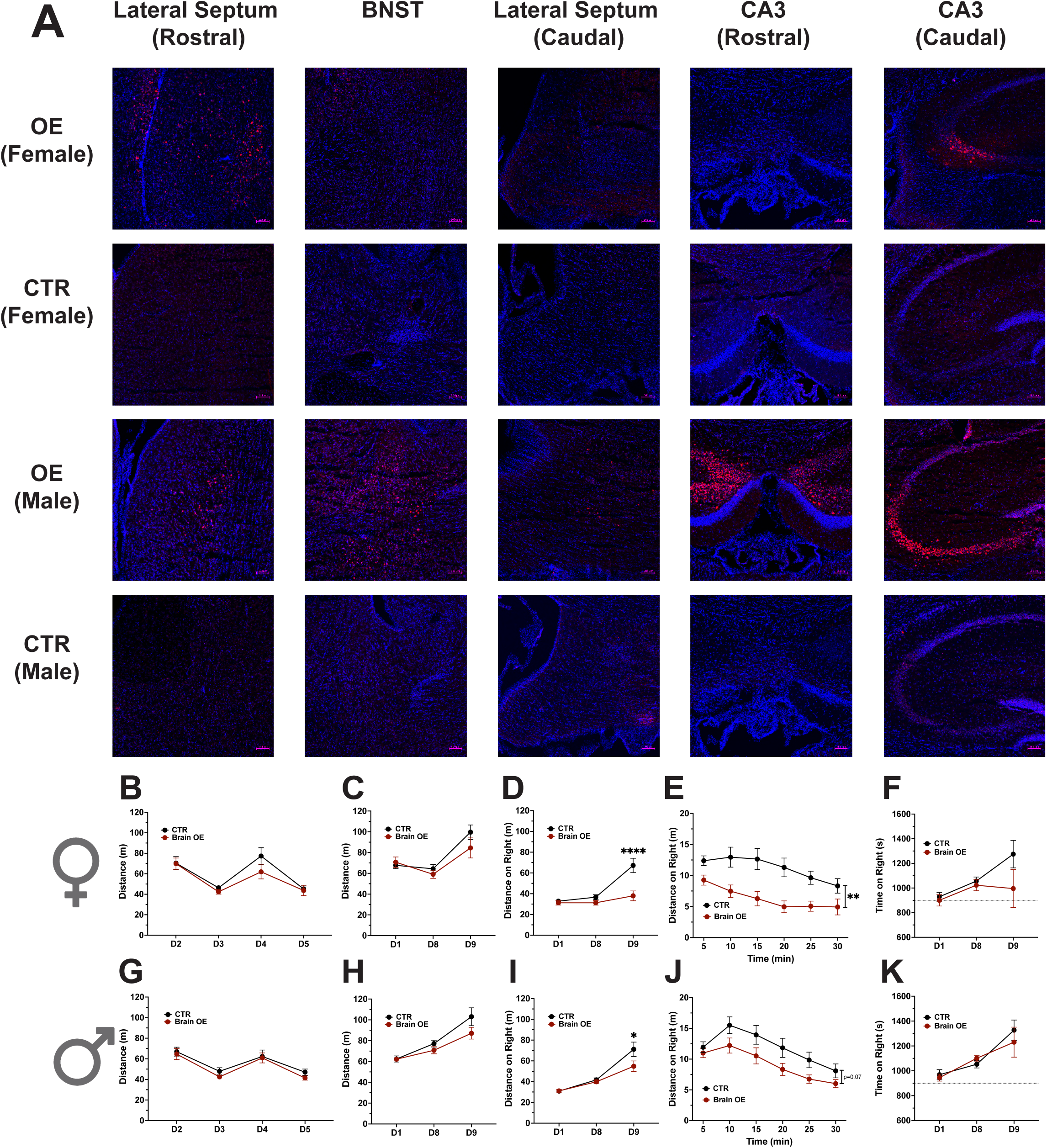
Effects of Zhx2 brain OE on brain OXY/metabolite levels and behavior in J females and J males following state-dependent OXY-CPP (1.25 mg/kg, i.p.). **(A):** Immunohistochemical staining in the LS, BNST, and CA3 following brain Zhx2 OE in J females and J males (Red: Zhx2; Blue: DAPI) **(B):** Total distance traveled on conditioning days (Days 2-5) in J females. There was a significant main effect of Day (F_3,90_ = 24.02, ****p < 0.0001) and no effect of AAV (F_1,30_ = 0.952, p = 0.347) or AAV x Day interaction (F_3,90_ = 1.316, p = 0.276). **(C):** Total distance traveled on testing days (Days 1, 8, 9) in J females. There was a significant main effect of Day (F_2,60_ = 9.76, **p < 0.01) and no effect of AAV (F_1,30_ = 0.354, p = 0.556) or AAV x Day interaction (F_2,60_ = 1.859, p = 0.176). **(D):** Total distance on right (OXY side) on testing days in J females. There was a significant effect of AAV (F_1,30_ = 12.37, p < 0.01**), Day (F_2,60_ = 14.57, ***p < 0.001) and a significant AAV x Day interaction (F_2,60_ = 9.33, **p < 0.01). Multiple comparisons indicated a significant reduction in total distance on right on Day 9 in OE J females (****p < 0.0001). **(E):** Total time on right on testing days in J females. There were no significant effects of AAV (F_1,30_ = 2.652, p = 0.114), Day (F_2,60_ = 3.781, p = 0.056), or AAV x Day interaction (F_2,60_ = 1.593, p = 0.218). **(F)**. Day 9 distance on right in J females across 5 min session bins. There was a significant effect of AAV (F_1,30_ = 12.29, **p < 0.01) and Time Bin (F_5,150_ = 10.09, ****p < 0.0001) but no significant AAV x Session Time interaction (F_5,150_ = 1.850, p = 0.161). **(G):** Total distance across conditioning days in J males. There was a significant main effect of Day (F_3,90_ = 26.24, ****p < 0.0001), but no effect of AAV (F_1,30_ = 0.657, p = 0.424) and no AAV x Day interaction (F_3,90_ = 0.246, p = 0.776). **(H):** Total distance across testing days in J males. There was a significant main effect of Day (F_2,58_ = 28.08, ****p < 0.0001) and AAV x Day interaction (F_2,58_ = 3.56, *p < 0.05). However, post-hoc analysis with correction for multiple comparisons revealed no significant differences between OE vs WT mice on any day. **(I):** Total distance on right across testing days in J males. There was a significant effect of Day (F_2,58_ = 38.77, ****p < 0.0001), and a significant AAV x Day interaction (F_2,58_ = 3.94, *p < 0.05). Multiple comparisons indicated a significant decrease in Day 9 distance on the OXY-paired side in OE J males (*p < 0.05). **(J):** Total time on right across testing days in J males. There was a significant effect of Day (F_2,58_ = 11.80, ***p < 0.001) but no effect of AAV (F_1,29_ = 1.077, p = 0.308) and no significant AAV x Day interaction (F_2,58_ = 0.161, p = 0.765). **(K):** Day 9 distance on right in J males across 5-min session bins. There was a significant effect of Time Bin (F_5,145_ = 34.07, ****p < 0.0001) but no significant effect of AAV (F_1,29_ = 3.55, p = 0.07) and no AAV x Time Bin interaction (F_5,145_ = 1.376, p = 0.258).

Brain Zhx2 OE in J females did not alter total distance across days (**Fig 2B-C**), but suppressed Day 9 distance on right (OXY-paired side; **Fig. 2D**) that was persistent across the entire 30 min session (**Fig. 2E**). While there was no significant change for time on right (**Fig. 2F**), time and distance on right were significantly correlated (R^2^ = 0.389; ****p<0.0001). As with J females, brain Zhx2 OE did not alter total distance across days in J males (**Fig. 2G-H**), but significantly decreased Day 9 distance on right (**Fig. 2I**), although this decrease was merely near significant when exclusively comparing within Day 9 across the 5 min session Time Bins (**Fig. 2J**). There was no effect of Zhx2 OE on time on right (**Fig. 2K**), but like females, there was a strong positive correlation between distance and time on right (R^2^ = 0.254; ***p < 0.001). Overall, Zhx2 overexpression in select brain regions lowered OXY state-dependent behavior in both sexes but with an outsized effect in females, generally consistent with our initial hypothesis.

### Effect of Zhx2 KO on brain OXY metabolite levels and behaviors in female and male By mice

Our results indicated that Zhx2 OE in the brain of J mice induced the predicted decrease in OXY behaviors. We wished to test whether *Zhx2* loss-of-function would be sufficient on the By background to increase brain OMOR concentration and OXY addiction-like behaviors via knocking out *Zhx2* exon 3 (the only coding exon; **E3KO**) in Bys (**Fig S3, Fig. 3A**). E3KO successfully ablated ZHX2 protein expression (**Fig 3B**). Consistent with our hypothesis, E3KO significantly increased brain OMOR levels in females (**Fig. 3C-E**), without significantly affecting OMOR in males (**Fig 3F-H**), thus recapitulating the QTL effect and providing strong support for *Zhx2* as a quantitative trait gene.

**Figure 3.**
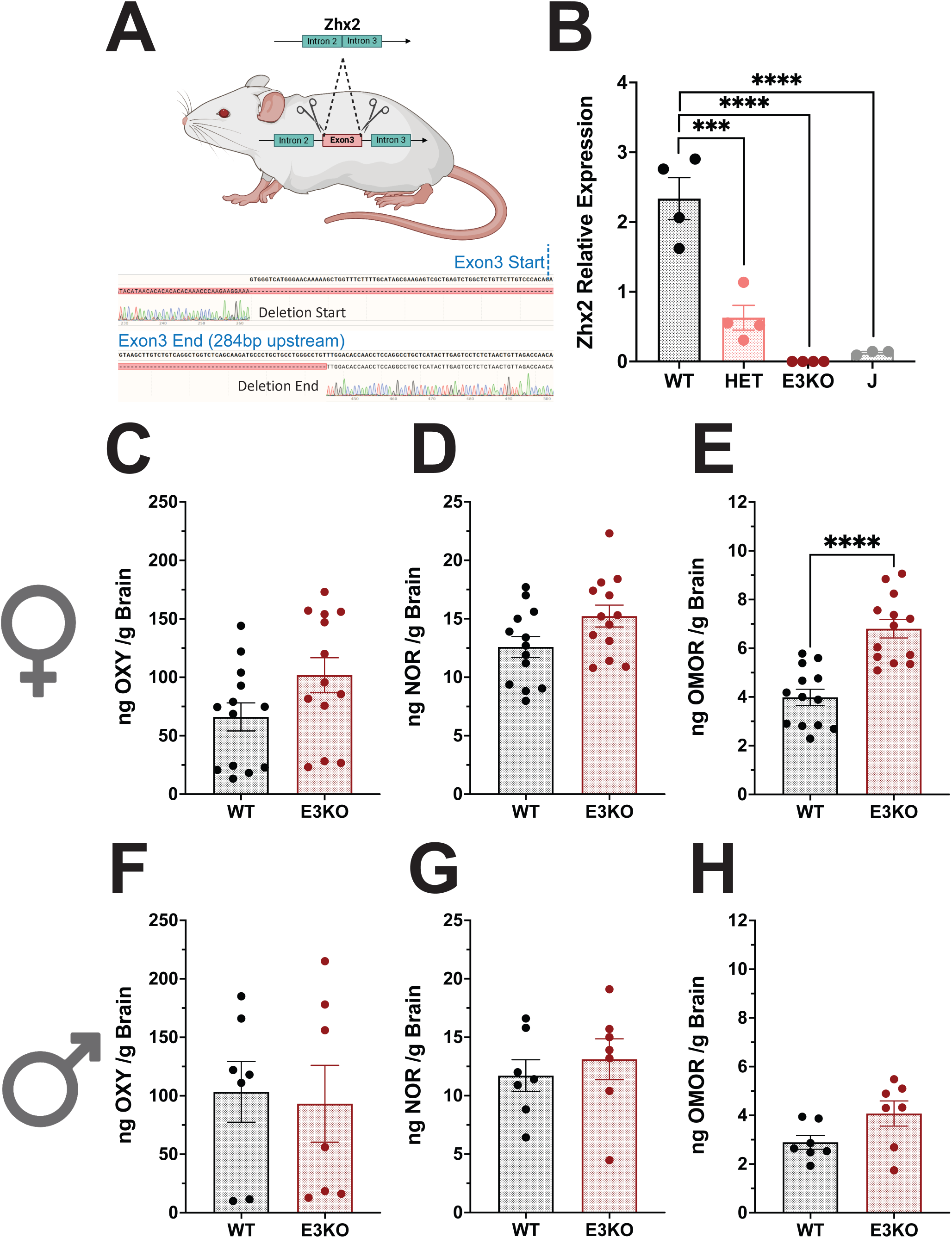
Brain OXY and metabolite quantification in Zhx2 knockout mice on BALB/cByJ (By) background following state-dependent OXY-CPP (1.25 mg/kg, i.p.). **(A):** Schematic of Zhx2 exon 3 deletion (E3KO) in Bys. Deletion began 78 bp upstream of Exon3’s proximal end and terminated 339 bp downstream of Exon3’s distal end. **(B):** Zhx2 protein quantification of all CRISPR offspring genotypes and BALB/cJs. There was a main effect of Genotype (F_3,11_ = 33.669, ****p < 0.0001) with multiple comparisons indicating significant differences between CRISPR WTs vs HET-E3KOs (***p < 0.001), WTs vs HOMO-E3KOs (****p < 0.0001), and WTs vs Js (****p < 0.0001). **(C):** Brain OXY concentrations between E3KO females versus WT By females (t_24_ = 1.86, p = 0.074). **(D):** Brain NOR concentrations in E3KO females versus WT females (t_24_ = 2.05, p = 0.051). **(E):** Brain OMOR concentrations in E3KO females versus WT By females (t_24_ = 5.58, ****p = 0.0001). **(F):** Brain OXY concentrations in E3KO males versus WT By males (t_12_ < 1). **(G):** Brain NOR concentrations in E3KO males versus WT By males (t_12_ < 1). **(H):** Brain OMOR concentrations in E3KO males versus WT By males (t_12_ = 2.01, p = 0.067).

E3KO females showed greater locomotor activity across conditioning days vs. WTs, and while the Genotype x Treatment x Day interaction was not significant, we observed OXY E3KO females traveling further than OXY WT females exclusively on Day 4 (second OXY exposure; **Fig. 4A**). E3KO females also traveled further across testing days vs. WTs (**Fig. 4B**). There was no significant effect of Genotype for distance on right or time on right across testing days in females (**Fig. 4C-D**). However, when examining change in time spent on the OXY-paired side between Day 9 vs. Day 1, time-dependent analysis revealed a significant reduction in OXY-CPP in E3KO females at the 10- and 15-min Time Bin (**Fig. 4E**), opposite of our prediction. In males, there were no significant effects of Genotype on total distance across all days (**Fig. 4F-G**), or in distance on right across testing days (**Fig 4H**). Surprisingly, we observed a significant Genotype x Treatment x Day interaction for time on right across testing days driven by a robust increase in OXY preference on Day 9 in OXY E3KO males (**Fig. 4I**), that was persistent across the 30 min session when comparing Day 9 vs. Day 1 across Time Bins (**Fig 4J**). In summary, *Zhx2* E3KO increased OXY locomotion and decreased OXY-CPP in females but increased state-dependent OXY-CPP without affecting OXY locomotion in males.

**Figure 4.**
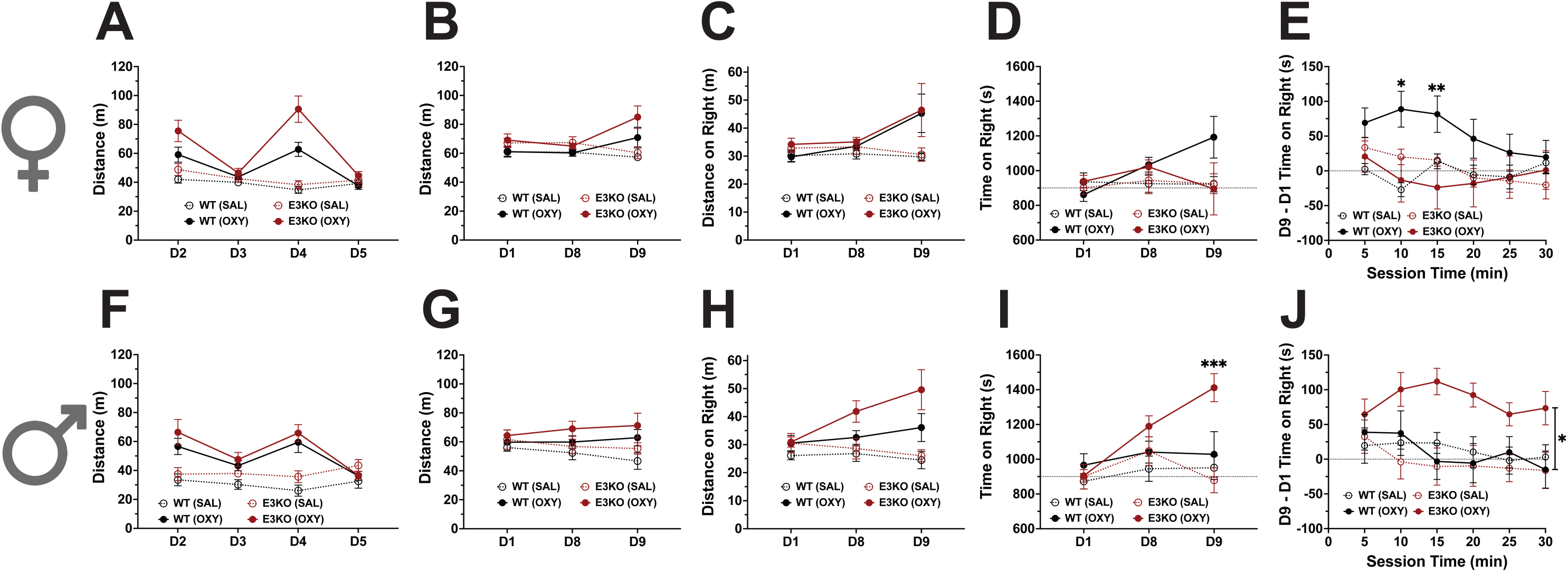
OXY locomotion and state-dependent OXY-CPP in Zhx2 E3KO mice in a BALB/cByJ (By) background. **(A):** Total distance traveled on conditioning days in females. There were significant effects of Genotype (F_1,52_ = 10.46, **p < 0.01), Treatment (F_1,52_ = 38.98, ****p<0.0001), Day (F_3,156_ = 19.74, ****p < 0.0001) and Treatment x Day (F_3,156_ = 22.84, ****p < 0.0001) However, there were no significant interactions involving Genotype x Treatment (F_1,52_ = 3.249, p = 0.077), Genotype x Day (F_3,156_ = 2.529, p = 0.072), or Genotype x Treatment x Day (F_3,156_ = 1.884, p = 0.135). Additionally, follow up analysis within each day yielded a significant Genotype x Treatment interaction on Day 4 (F_1,52_ = 4.767, *p < 0.05) with no significant Genotype x Treatment interactions on Day 2 (F_1,52_ = 0.862, p = 0.357), Day 3 (F_1,52_ = 0.001, p = 0.970), or Day 5 (F_1,52_ = 0.555, p = 0.460). Multiple comparisons on Day 4 revealed that OXY E3KO females traveled significantly further than OXY WT females (t_52_ = 3.611, **p < 0.01) with no significant differences in distance traveled between SAL E3KO females and SAL WT females (t_52_ < 1, p = 0.654). **(B):** Total distance traveled on testing days in females. There were significant effects of Genotype (F_1,53_ = 8.08, **p<0.01), Treatment (F_1,53_ = 6.67, *p < 0.05), and Treatment x Day (F_2,106_ = 6.29, **p < 0.01). However, there was no significant effect of Day (F_2,106_ = 1.531, p = 0.223) and no significant interactions involving Genotype x Treatment (F_1,53_ = 2.267, p = 0.138), Genotype x Day (F_2,106_ = 0.184, p = 0.810), or Genotype x Treatment x Day (F_2,106_ = 0.274, p = 0.738). **(C):** Total distance on right traveled on testing days in females. There was a significant effect of Treatment (F_1,53_ = 5.27, *p < 0.05) and a significant Treatment x Day interaction (F_2,106_ = 5.46, *p < 0.05). However, there was no effect of Genotype (F_1,53_ = 1.68, p = 0.200), or Day (F_2,106_ = 2.673, p = 0.096), and no significant interactions involving Genotype x Treatment (F_1,53_ = 0.368, p = 0.547), Genotype x Day (F_2, 106_ = 0.210, p = 0.720), or Genotype x Treatment x Day (F_2,106_ = 0.154, p = 0.768). **(D)**. Total time on right traveled on testing days in females. There was a significant effect of Day (F_2,106_ = 3.57, *p < 0.05) but no significant effect of Genotype (F_1,53_ = 0.159, p = 0.691) or Treatment (F_1,53_ = 0.435, p = 0.512), and no significant interactions involving Genotype x Treatment (F_1,53_ = 0.459, p = 0.501), Genotype x Day (F_2,106_ = 3.565, p = 0.093), Treatment x Day (F_2,106_ = 1.229, p = 0.289), or Genotype x Treatment x Day (F_2,106_ = 2.282, p = 0.122). **(E)**. Time on right on Day 9 – Day 1 across 5 min session time bins. There was a significant effect of Time Bin (F_5,265_ = 3.25, p < 0.01**) and a Genotype x Treatment x Time Bin interaction (F_5,265_ = 3.98, **p < 0.01). There was no significant effect of Genotype (F_1,53_ = 2.513, p = 0.119) or Treatment (F_1,53_ = 1.721, p = 0.195) and no significant interactions involving Genotype x Treatment (F_1,53_ = 3.780, p = 0.057), Genotype x Time Bin (F_5,265_ = 1.015, p = 0.394), or Treatment x Time Bin (F_5,265_ = 0.487, p = 0.718). Multiple comparisons revealed a significant decrease in OXY-CPP in E3KO females at 10 min (*p < 0.05) and 15 min (**p < 0.01). **(F):** Total distance traveled on conditioning days in males. There were a significant effects of Treatment (F_1,32_ = 21.06, ****p < 0.0001) and Day (F_3,96_ = 6.15, **p < 0.01), as well as a significant Treatment x Day interaction (F_3,96_ = 11.83, ****p < 0.0001). However, there was no effect of Genotype (F_1,32_ = 3.237, p = 0.081) and no significant interactions involving Genotype x Treatment (F_1,32_ = 0.147, p = 0.704), Genotype x Day (F_3.96_ = 0.058, p = 0.955) or Genotype x Treatment x Day (F_3.96_ = 0.590, p = 0.573). **(G):** Total distance traveled on testing days in males. There was a significant effect of Treatment (F_1,31_ = 5.34, *p < 0.05). However, there were no significant effects of Genotype (F_1,31_ = 0.958, p = 0.335) or Day (F_2,62_ = 0.101, p = 0.894), and no significant interactions involving Genotype x Treatment (F_1,31_ = 0.621, p = 0.437), Genotype x Day (F_2,62_ = 1.521, p = 0.228), Treatment x Day (F_2,62_ = 1.961, p = 0.149), or Genotype x Treatment x Day (F_2,62_ = 0.668, p = 0.509). **(H):** Total distance on right traveled on testing days in males. There was a significant effect of Treatment (F_1,31_ = 14.38, ***p < 0.001) and a significant Treatment x Day interaction (F_2,62_ = 4.97, *p < 0.05). However, there were no significant effects of Genotype (F_1,31_ = 0.882, p = 0.355) or Day (F_2,62_ = 1.555, p = 0.223), and no significant interactions involving Genotype x Treatment (F_1,31_ = 0.038, p = 0.846), Genotype x Day (F_2,62_ = 2.398, p = 0.112), or Genotype x Treatment x Day (F_2,62_ = 2.505, p = 0.103). **(I)**: Total time on right on testing days in males. There was a significant effect of Treatment (F_1,31_ = 16.41, ***p < 0.001), and Day (F_2,62_ = 4.66, *p < 0.05), as well as significant interactions involving Treatment x Day (F_2,62_ = 3.32, *p < 0.05), and Genotype x Treatment x Day (F_2,62_ = 5.29, **p < 0.01). However, there was no significant effect of Genotype (F_1,31_ = 0.700, p = 0.409) and no significant interactions involving Genotype x Treatment (F_1,31_ = 0.776, p = 0.385) or Genotype x Day (F_2,62_ = 2.157, p = 0.130). Multiple comparisons revealed a significant increase in state-dependent OXY-CPP in E3KO males versus WT males on Day 9 (***p < 0.001). **(J)**: Time on right on Day 9 – Day 1 across 5 min Time Bins in males. There was a significant effect of Treatment (F_1,31_ = 5.69, **p < 0.05) and a significant Genotype x Treatment interaction (F_1,31_ = 6.40, *p < 0.05). However, there was no significant effect of Genotype (F_1,31_ = 2.589, p = 0.118) or Time Bin (F_5,155_ = 1.902, p = 0.127), and no significant interactions involving Genotype x Time Bin (F_5,155_ = 0.357, p = 0.806), Treatment x Time Bin (F_5,155_ = 0.428, p = 0.755), or Genotype x Treatment x Time Bin (F_5,155_ = 1.843, p = 0.137). Subsequent follow-up indicated an overall difference between OXY treated WTs vs. OXY treated E3KOs across the session (F_1,17_ = 8.45, *p<0.05).

### Multi-omics analysis of Zhx2 E3KO in the brain

While the female-specific E3KO increase in OMOR supports validation of *Zhx2* as a quantitative trait gene[32], the behavioral results, while supporting *Zhx2’s* sex-dependent influence on OXY addiction-like behaviors, were surprising and could indicate additional mechanisms beyond metabolism that connect *Zhx2* loss-of-function with OXY addiction-like behaviors. Thus, we performed exploratory multi-omics analysis in E3KOs. Complete lists of genes/proteins and enriched pathways are provided in **Tables S3-20**. Top differentially expressed genes and proteins by sex are annotated in **Fig. S11,** which included the related ZHX1 and ZHX3 in addition to multiple astrocyte markers (Gfap, ALDH1L1, ETNPPL)[57–59]. Combining -omics datasets within each sex for gene set enrichment analysis yielded top shared enrichment terms, including extracellular matrix organization, cell adhesion, inflammatory responses, oxoacid metabolic processes, and carboxylic acid catabolic processes (**Fig. 5A-B**). Top differentially expressed genes/proteins exclusive to one sex (p.adj < 0.05 in one sex, p.unadj > 0.05 in opposite sex) included the brain endothelial cell marker SLCO1C1 in females and the oxidative stress protein PON2 in males (**Fig. S12A-B**). Enriched pathways from omics-combined differentially expressed genes/proteins exclusive to one sex (p.unadj < 0.05 in one sex, p.unadj > 0.05 in opposite sex) yielded regulation of extracellular matrix organization in females and glial cell development in males (**Fig. S12C-D**).

**Figure 5.**
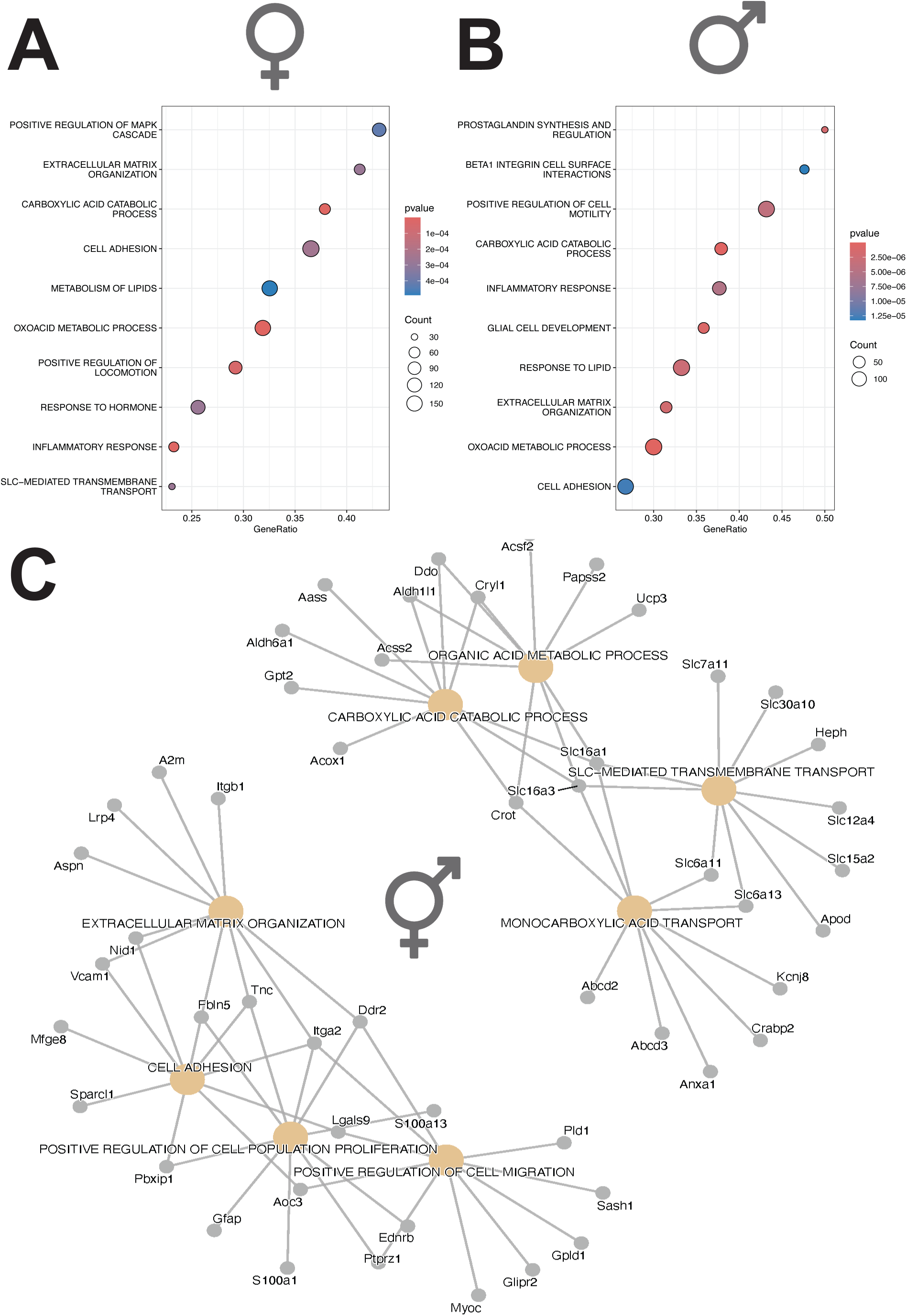
Zhx2 E3KO -Omics Pathway Enrichment. **(A):** Top 10 enriched pathways in E3KO females using Gene Set Enrichment Analysis on combined transcriptomics/proteomics p-value rank ordered differentially expressed genes/proteins. Pathways ordered by leading-edge Gene Ratios. **(B):** Top 10 enriched pathways in E3KO males using Gene Set Enrichment Analysis on combined transcriptomics/proteomics p-value rank ordered differentially expressed genes/proteins. Pathways are ordered by leading-edge Gene Ratios. **(C):** Cnetplot of top 8 enriched pathways in sex-collapsed E3KOs vs WTs using Gene Set Enrichment Analysis on combined transcriptomics/proteomics p-value rank ordered differentially expressed molecules. Interactions are shown between top 10 leading-edge genes per pathway and pathways they are a part of.

To assess pathway similarity and common genetic factors, we collapsed across sex and -omics datasets and visualized the top 8 enriched pathways with the top 10 leading-edge molecules for each pathway (**Fig. 5C**). Key molecules contributing to at least 3 pathways included TNC, FLBN5, ITGA2, and DDR2, which are all important for extracellular matrix function and cell adhesion[60–63], SLC16A3 and SLC16A1, which are lactate transporters associated with astrocyte and endothelial cell expression and activity[64–67], and CROT, which can influence fatty acid metabolism and mitochondrial function[68]. Using STRING-db to visualize interactions between these leading-edge molecules indicated GFAP expression associated with multiple cell clusters (**Fig. S13,** full cluster results in **Table. S21**). Overall, our results suggest *Zhx2* loss-of-function could compromise astrocyte function which in turn could disrupt multiple biological processes (extracellular matrix function, endothelial cell activity, and mitochondrial dysfunction) with a sex-specific bias in certain pathways.

## DISCUSSION

The major goal of this study was to test for validation of *Zhx2* as a quantitative trait gene underlying brain OMOR concentration associated with OXY addiction-like behavior. While constitutive germline restoration of *Zhx2* function via MERV removal was insufficient to reduce brain OMOR or alter OXY behavior on the mutant J genetic background, Zhx2 OE in the brain vs. liver provided support for *Zhx2* function in the brain in mediating OXY behavior. Additionally, *Zhx2* loss-of-function on the wild-type By genetic background was sufficient to increase brain OMOR concentration and modulate OXY behavior in a sex-specific manner, providing support for *Zhx2* as a quantitative trait gene[32]. Furthermore, we identified key molecular adaptations by which *Zhx2* could regulate OXY metabolite levels and behavior, including astrocyte activity, cell adhesion, inflammation, cellular function and oxidative stress, and extracellular matrix function.

The lack of effect of liver OE on brain OXY and metabolite concentrations was unexpected, considering sex differences in liver Cyp2d22 RNA expression, a result complementing previous work detailing that hepatic Zhx2 expression influences sex-specific liver Cyp expression[37]. While the function of individual mouse CYP2D enzymes remain largely unexplored, CYP2D22 was identified as the closest ortholog to the human CYP2D6 in structure and opioid-metabolizing function[56,69], and thereby could metabolize OXY to OMOR. However, the lack of brain metabolite differences in response to 100-fold increase in liver Zhx2 expression indicates that if *Zhx2* regulates brain OMOR levels through CYP2D enzymes, it is likely to do so primarily in the brain. Alternatively, the massive overexpression could engage additional mechanisms that obfuscate potential brain OMOR changes caused by milder, biologically relevant increases in Zhx2 expression.

While *Zhx2* OE was not widespread throughout the brain, there was decreased OXY state-dependent behavior that was associated with OE in the LS, BNST, and CA3 of the hippocampus. AAV i.c.v. injections with other serotypes (AAV9 and AAV.PHP.B) in BALB/c mice also did not spread across the whole brain[70]. We chose the AAV/F serotype based on its ability to more effectively transfect the whole brain compared to AAV9 through multiple administration routes and because, unlike AAV.PHP.B, AAV/F transfects the whole brain to a similar extent between C57BL/6 and BALB/c strains when administered systemically[71]. However, a direct comparison between whole brain viral transfection in C57BL/6 vs. BALB/c using the i.c.v. route was not reported. Nonetheless, localized OE in these brain regions was sufficient to decrease OXY state-dependent behaviors. The LS contributes to motivational behaviors and cue-reward conditioning[72,73]. The BNST is widely implicated in drug related behaviors, especially withdrawal[74–76]. CA3 suppression is shown to impair overall spatial learning, memory, and drug environmental context-induced reinstatement[77–79]. Therefore, these regions could influence opioid state-dependent learning and environmental conditioning.

As predicted based on our QTL study, *Zhx2* loss-of-function via E3KO increased brain OMOR in females but not males, supporting *Zhx2* as the causal gene underlying increased brain OMOR concentrations in females inheriting the *Zhx2* MERV loss-of-function[32]. While we were surprised that ZHX2 expression restoration via MVKO did not reciprocally lower brain OMOR in females, one potential reason may involve epistatic interactions with *Zhx2* that depend on the J vs. By genetic background, thereby influencing whether manipulating ZHX2 expression can alter downstream phenotypes. Additionally, differences in environmental exposure from mice bred in house (Zhx2 CRISPR lines) versus ordered from JAX (for Zhx2 tissue-specific OE studies) could influence behavior[80,81]. This could also explain why Js ordered from JAX exhibited robust OXY state-dependent learning[32], while MVKO WT J mice bred in house did not. Furthermore, indirect social genetic effects on behavior based on different genotypes in cage moms and/or littermates could also explain these differences[82,83].

Behaviorally, observed locomotor increases in female E3KOs complement our previous work illustrating greater OXY-induced locomotion in Js, seemingly driven by females[32]. While E3KO altered OXY state-dependent learning in a sex dependent manner, neither the presence of state-dependent learning in both sexes or the directionality was in line with our original hypothesis. The original QTL was for brain OMOR concentration, as we were not powered to detect QTLs for state-dependent OXY-CPP behaviors[32]. Thus, substrain differences in state-dependent OXY behaviors could be regulated by additional genetic differences, including any one of the nine Cyp2d genes that also reside in the QTL interval which harbors *Zhx2*[32]. Notably, the state-dependent OXY-CPP reduction in E3KO females was localized at 10-15 min post-OXY, the expected Tmax brain OXY concentration following i.p. administration in rodents[84]. While sex comparisons in OXY pharmacokinetics are limited, one study reported higher initial and max OXY concentrations after multiple administration routes in female rodents[85], suggesting sex-dependent OXY pharmacokinetic differences could contribute to behavioral differences.

Suppressing ZHX2 in Bys yielded no detectable differences in brain Cyp2d enzyme expression, likely due to overall low Cyp2d brain expression, and thereby may require increased sequencing and fractionation depth to test for differential expression between genotypes. However, we observed several enriched pathways related to lipid metabolism, immune function, and cellular processes, all in line with established roles for *Zhx2*[36]. Noteworthily, multiple astrocyte markers were differentially expressed, as *Zhx2* is predominantly expressed in astrocytes[86]. Visualizing interactions between top leading-edge genes yielded interactions between GFAP and several clusters, indicating *Zhx2* could alter astrocyte function to exert widespread consequences on brain function. Accordingly, we observed enrichment of extracellular matrix and cell adhesion pathways. Accumulating evidence indicates extracellular matrix activity can influence opioid behavior via neuroinflammation and synaptic signaling[87]. Discoidin domain receptor tyrosine kinase II (**DDR2**), both a leading edge molecule regulating multiple top pathways and a top differentially expressed molecule across multiple data sets, is associated with multiple alcohol and nicotine behavioral outcomes[88–90]. Knocking out DDR2 in cells lowers matrix metalloproteinase (**MMP**) expression and cell motility[60]. MMP activity is associated with opioid misuse[91,92] and thus could explain how *Zhx2* manipulations effect opioid behavioral outcomes. While the inflammatory response pathway was upregulated in both sexes, multiple inflammatory molecules were lowly expressed or undetectable, possibly because our -omics experiments were solely performed in OXY-naive mice. Therefore, it would be useful for future work to assess how OXY treatment interacts with *Zhx2* to influence these pathways, including OXY-induced neuroinflammation[93].

Paraoxonase 2 (**PON2**) was the top differentially expressed molecule in males, being upregulated in KOs. PON2 is commonly associated with oxidative stress and inflammation, is expressed in dopaminergic reward related areas like the striatum, is expressed predominately in astrocytes more than neurons, and is associated with smoking outcomes[95,96]. As we observed multiple upregulated pathways related to oxidative stress, ZHX2 ablation could increase OXY state-dependent learning in males via altered PON2 expression. SLCO1C1 was a top downregulated protein in females. Solute carrier organic anion transporter family member 1C1 (**SLCO1C1**), a thyroid hormone transporter at the blood brain barrier **(BBB)**[97], is specific to brain endothelial cells and can alter BBB functionality[98–100]. The BBB is composed of endothelial cells and astrocytic end feet to form tight cell adhesions to protect neural tissues[101,102], and its integrity depends critically on extracellular matrix activity[103]. The BBB regulates entry of opioids and thus is critical to influencing their addictive properties[104]. As we observed multiple enriched pathways relating to astrocyte function, cell adhesion, extracellular matrix activity, and endothelial cells, *Zhx2* could influence BBB function to increase OXY and/or OXY metabolite and behavioral changes in females.

Overall, our results illustrate *Zhx2* can regulate OXY metabolite levels and OXY addiction-like behaviors in a sex-dependent manner. As *Zhx2* was previously associated with nicotine use in humans[69,70], these observations combined with our previous work[32] provide further evidence that *Zhx2* can influence addiction-like traits for substance use disorders. Further assessment of identified downstream mechanisms could uncover novel therapeutic targets to aid novel individually tailored OUD treatments.

## Supporting information

Supplemental Table 3. RNA-seq gene list (Genotype - Sex covariate).

Supplemental Table 4. RNA-seq gene list (Genotype - Males).

Supplemental Table 5. RNA-seq gene list (Genotype - Females).

Supplemental Table 6. Mass-spectrometry protein list (Genotype - Sex covariate).

Supplemental Table 7. Mass-spectrometry protein list (Genotype - Males).

Supplemental Table 8. Mass-spectrometry protein list (Genotype - Females).

Supplemental Table 9. Multi-omics gene/protein list (Genotype - Sex covariate).

Supplemental Table 10. Multi-omics gene/protein list (Genotype - Males).

Supplemental Table 11. Multi-omics gene/protein list (Genotype - Females).

Supplemental Table 12. RNA-seq pathway enrichment list (Genotype - Sex covariate).

Supplemental Table 13. RNA-seq pathway enrichment list (Genotype - Males).

Supplemental Table 14. RNA-seq pathway enrichment list (Genotype - Females).

Supplemental Table 15. Mass-spectrometry pathway enrichment list (Genotype - Sex covariate).

Supplemental Table 16. Mass-spectrometry pathway enrichment list (Genotype - Males).

Supplemental Table 17. Mass-spectrometry pathway enrichment list (Genotype - Females).

Supplemental Table 18. Multi-omics pathway enrichment list (Genotype - Sex covariate).

Supplemental Table 19. Multi-omics pathway enrichment list (Genotype - Males).

Supplemental Table 20. Multi-omics pathway enrichment list (Genotype - Females).

Supplemental Table 21. MCL clustering of top leading-edge gene/proteins.

## ACKNOWLEDGMENTS

We thank the Institute for Chemical Imaging of Living Systems (RRID:SCR_022681) at Northeastern University for consultation and imaging support. We also thank The University of Chicago Genomics Facility (RRID:SCR_019196) especially Pieter Faber, Technical Director for the University of Chicago Genomics Facility, for their assistance with the Illumina NovaSeqX. We also thank the National Institutes of Health for support in funding the Orbitrap Eclipse Tribrid Mass Spectrometer (S10OD026807) used in this work.

## AUTHOR CONTRIBUTIONS

W.B.L.: Experimental design, data collection, analysis, and writing of manuscript; S.A.M: Data collection and analysis; S.I.G: Experimental design, data collection, analysis, and writing of manuscript; J.A.B.: Experimental design; R.B.: Data collection and analysis; E.T.G.: Data collection and analysis; A.F.: data collection and analysis; B.N.: Data collection and analysis; K.K.W.: Data collection; I.K.: Data collection and analysis; G.A.S.: Data collection and analysis; O.A.: Data collection and writing of manuscript; B.M.B.: Data collection; M.T.F.: Experimental design; C.A.R.: Data collection and writing of manuscript; A.E.: Data collection and writing of manuscript; C.D.B.: Experimental design, data collection, analysis, and writing of manuscript.

## FUNDING

This work was supported by a NIDA-F31-DA056217 to WBL, a NIDA-T32-DA055553 to BMB and a NIDA-U01-DA050243 to CDB

## COMPETING INTERESTS

The authors have nothing to disclose.

## DATA AVAILABILITY STATEMENT

The RNA-sequencing datasets generated and analyzed during the current study are available in the Gene Expression Omnibus repository (GSE275965). All other datasets generated during and/or analyzed during the current study are available from the corresponding author on reasonable request.

## Notes

### Competing Interest Statement

The authors have declared no competing interest.

