## Supplemental Materials for "Validation studies and multi-omics analysis of Zhx2 as a candidate quantitative trait gene underlying brain oxycodone metabolite (oxymorphone) levels and behavior"

#### SUPPLEMENTARY MATERIALS AND METHODS

##### Mouse Breeding and Genotyping

Embryos were injected with guide RNAs to direct the Cas9 protein towards relevant sites: *Zhx2* MERV in BALB/cJs (JAX: 000651) and *Zhx2* Exon3 in BALB/cByJs (JAX: 001026). The 5.6 kb *Zhx2* MERV insertion was removed from the J background via CRISPR-mediated excision and replaced with the wild-type By sequence using a homologous, complementary oligonucleotide. CRISPR-deletion in Js began directly at the MERV's proximal end and terminated directly at the MERV's distal end within intron 1[1,2]. Reciprocally, the 5.6 kb MERV Zhx2 loss-of-function was modeled on the By background via knockout of exon 3, the sole coding exon (inserting the 5.6 kb MERV into the By genome could not be achieved by JAX for reasons suspected to be related to the BALB/c genome). CRISPR-deletion in Bys began 78 bp upstream of the proximal end and terminated 339 bp downstream of the distal end of exon 3. Founders (**F1**) were backcrossed to their respective parental strain before resultant heterozygous offspring (**N1**) were sent to the Bryant lab for subsequent breeding. Upon arrival to the Bryant lab, offspring were backcrossed again to their parental strains of origin with subsequent heterozygous offspring crossed to generate wild-type (**WT**), heterozygous, and homozygous knockout mice. WTs and either MERV knockouts (**MVKOs**) or Exon3 knockouts (**E3KOs**) were used for subsequent experiments.

Tail snips were taken at weaning (21 days post-natal) for subsequent genotyping. DNA was extracted and we assessed the presence vs. absence of the MERV in J CRISPR offspring and exon 3 in By CRISPR offspring. Sequences were amplified using DreamTaq MasterMix (ThermoFisher, Waltham, MA) with a PCR Thermal Cycler T100 (Bio-Rad, Hercules, CA). Amplified sequences were visualized using a 2% agarose gel with ethidium bromide.

##### AAV Administration

For liver AAV targeting, mice were briefly anesthetized with 4% isoflurane and placed in a nose cone with 2% isoflurane during administration. One drop of 0.5% proparacaine hydrochloride (Sandoz, Basel, Switzerland) was placed on the eye 15 s prior to AAV administration for analgesia. The following AAVs were administered retro-orbitally for liver Zhx2 overexpression **(OE)**: AAV8-TBG-mZhx2-P2A-eGFP (OE), AAV8-TBG-eGFP (Control; **CTR**), with administration behind one eye into the medial canthus using a 27-gauge, 0.5 inch insulin needle[3]. The needle was held in place for 2 seconds before slowly withdrawn to ensure distribution. Each injection contained a $1x10^{11}$ gc/mouse viral load and a 100ul volume[4,5].

For brain AAV targeting, mice were anesthetized with 4% isoflurane and head fixed in a stereotaxic apparatus. Mice were closely monitored throughout the surgery with anesthesia maintained at 1-2% isoflurane. The lateral ventricle (**LV**) was targeted using the following coordinates relative to bregma (in mm): anterior/posterior (A/P): -0.30, medial/lateral (M/L): +1.00, dorsal/ventral (D/V) -2.60 (from the dura)[6]: The following AAVs were administered intracerebroventricularly **(i.c.v.)** for brain Zhx2 OE: AAV/F-CMV-mZhx2-P2A-eGFP.miR122 (OE), AAV/F-CMV-eGFP.miR122 (CTR). Virus was delivered unilaterally to the left LV with a 33-gauge needle attached to a 5ul syringe (Hamilton, Reno, NV). Virus was delivered at a rate of 1ul every 30 sec for 5 min for a total infusion volume of 10ul, with a $5x10^{10}$ gc/mouse viral load[7]. The needle was left in place for an additional 5 min to facilitate/ensure distribution into the LV.

**Conditioned Place Preference**

All sessions were 30 min long and all animals either received OXY (1.25mg/kg; 0.01ml/kg; i.p.) based on daily body weight or an equivalent volume of saline (**SAL;** 0.01ml/kg; i.p.). On Day 1 mice ages 50-100 days old were injected with SAL and placed on the left side (smooth floor texture) with open access to both chambers. On Days 2 & 4 mice received OXY (or SAL control) and were confined to right side of the chamber (pointed floor texture) while on Days 3 & 5, all mice received SAL and were confined to the left side (again, smooth floor texture). On Day 8 mice were again injected with SAL, placed on the left side, and had open access to both sides to assess CPP. On Day 9, mice were injected with OXY (or SAL for controls), placed on the left side, and had open access to assess state-dependent learning of CPP. All sessions were video recorded using infrared cameras (Swan, Victoria, Australia) and tracked using ANY-Maze (Stoelting, Wood Dale, IL).

**Immunohistochemistry**

Brains were embedded in O.C.T. Compound (ThermoFisher), and kept at -20degC. Coronal slices (30 um thickness) were collected on a cryostat and mounted directly onto glass slides. The whole brain (prefrontal cortex to spinal cord) was sliced and collected to locate Zhx2 expression. Slides were stored in -80degC until further processing. For immunostaining, slides were re-hydrated in 3x5 min 1X phosphate-buffered saline (**PBS**) washes, then incubated in blocking solution (5% normal donkey serum, 0.2% Triton X-100, and 1X PBS) for 1 hr at room temperature, then incubated in primary anti-ZHX2 antibody (GeneTex; 1:750) at 4degC overnight. Tissue was washed in 1X PBS (3x5 mins) then incubated in donkey anti-rabbit secondary antibody (ThermoFisher; 1:750) at room temperature for 1 hr. Slides were washed in 1X PBS (3x5 min) and incubated in DAPI (ThermoFisher; 1:100) solution for 5 min. Tissue was cover slipped with VectaShield mounting medium (Vector Laboratories, Newark, CA) and dried at room temperature. Fluorescence was visualized on a Zeiss LSM 800 Confocal Microscope (Zeiss, Danvers, MA) with the same brightness and exposure on each image. Slides were stored at 4degC for long-term storage. Images were processed using ImageJ[8] with changes in brightness and contrast applied evenly across all images.

**Proteomic Mass-Spectrometry**

Proteomic measurements and analyses were performed in flash-frozen left-brain hemispheres in SAL-treated mice. Brains were homogenized and proteins were extracted, digested with trypsin, desalted, and labelled with TMTpro reagents (ThermoFisher) as described previously[9]. The labelled peptides were fractionated using high pH reversed-phase chromatography on an Agilent 1260 Infinity Capillary LC equipped with an XBridge Peptide BEH C18 column (Waters Corperation, Milford, MA). Pooled fractions were separated on an Easy-nLC 1200 system connected to an Orbitrap Eclipse mass spectrometer equipped with a FAIMS Pro Interface (ThermoFisher). MS/MS spectra were searched using the Andromeda search engine in

MaxQuant software (version 2.0.3.0) against the complete SwissProt mouse proteome and the MaxQuant contaminants database[10,11]. Peptide spectrum match and protein FDRs were set at 0.01 using the target-decoy database search strategy[12].

**Multi-Omics Pathway Enrichment and Visualization**

Pathway enrichment analysis was performed on the transcriptomic, proteomic, and combined multi-omic list of differentially expressed genes and proteins (see Statistical Analyses regarding details of combined -omics analysis). Analysis was performed comparing *Zhx2* E3KOs vs. WTs within each sex separately and with sex as an additive co-variate (n=4 per Genotype per Sex). Pathways were obtained from a curated list across multiple different sources[13]. Background genes/proteins were set to only include genes/proteins within our data sets. Gene/protein sets were ranked by the absolute log10 p-values. Pathway enrichment analysis was performed on subsequent gene/protein rankings using the fgsea R package with an adaptive multilevel splitting approach and a minimum pathway size set to 15 genes and maximum set to 500 genes[14,15]. The collapsePathways command in fgsea was used to remove redundant/similar pathways that were significant ($p < 0.05$).

Log2FC was calculated for each gene and protein and visualized using the EnhancedVolcano R package[16]. Genes and proteins with a p-value below 0.05 and a logFC above 0.2 in E3KOs vs WTs were considered differentially expressed, with top differentially expressed genes and proteins annotated. The top 10 enriched pathways by p-value from the multi-omics ranked lists within each sex were plotted using the dotplot function in the enrichplot R package[17]. Enriched pathways were listed by gene ratio (leading-edge genes/total genes in pathway) with circle size indicating number of leading-edge genes within each pathway and color indicating the pathway p-value. Pathway interactions were visualized using the cnetplot function in enrichplot. The top 8 significant pathways using the multi-omic sex combined gene/protein rankings were visualized along with the top 10 leading-edge genes of each pathway. Each grey circle represents a leading-edge gene with yellow circles representing pathways, with lines connecting circles indicating a leading-edge gene contributes towards a respective pathway(s). Interactions between these unique top 10 leading-edge genes from these top 8 pathways were visualized using STRING-db[18]. Resultant genes were clustered using a Markov Cluster Algorithm[19], with node color indicating the associated cluster. Interactions between genes were indicated by connecting lines, with line thickness indicating interaction strength based upon pre-existing sources. Solid lines indicate interactions within a cluster while dotted lines indicate gene interactions from different clusters.

Unique enriched pathways for each sex were determined using Metascape from combined gene/protein lists significant in one sex but not the other (p.unadj < 0.05 vs. p.unadj > 0.05), done for both females and males[20]. The top 20 enriched pathways for each sex were ranked by enrichment p-value and plotted.

**Statistical Analyses**

Metabolite and qPCR data were analyzed with unpaired student's t-tests. A Welch's correction was applied for any group that did not have equal variances as determined by an F-test of equal variances. ZHX2 immunoblots were analyzed using one-way ANOVAs. Behavioral data summed across a 30 min session across days or data broken down into 5 min session intervals within each day were analyzed with mixed-effect ANOVAs with repeated measures for Day or Time Bin. Sphericity of the data was assessed using Machly's Test for Sphericity with an alpha level of 0.05. If sphericity was violated ($p < 0.05$), a Greenhouse-Geisser correction was applied toward ANOVA p-values for within-subject main effects or interactions. All statistics were performed separately in females and males due to our leading hypothesis of female specific phenotypic changes[21], with the exception of Zhx2 RNA or protein quantification for target validation purposes.

For a significant main effect of Genotype in ZHX2 western blotting, follow-up multiple comparisons were performed between all Genotype levels. For significant 3-way interactions between Genotype, Treatment, and either Day or Time Bin, follow-up multiple comparisons were performed by comparing Genotype differences within each treatment group (SAL or OXY) and within each Day or 5 min Time Bin. For 2-way interactions between Genotype and Treatment without a Day or Time Bin interaction, follow-up multiple comparisons were performed comparing Genotype differences within each treatment (SAL or OXY), across Day or 5 min Time Bin. P-values were generated via pair-wise comparisons from the estimated marginal means then adjusted using the Holm-Sidak correction[22]. Significance was defined as follows: *$p < 0.05$, **$p < 0.01$, ***$p < 0.001$, ****$p < 0.0001$.

For transcriptomic differential gene expression, data were normalized using a trimmed mean of M values (**TMM**) between sample pairs before using the edgeR quasi-likelihood pipeline for differential gene expression[23]. The error rate for multiple testing was controlled using Benjamini and Hochberg's false discovery rate set at a 0.05 threshold. For proteomic analysis, protein intensities were log transformed and Loess normalized. Linear models were fitted to each gene and group comparisons between factors were conducted using moderated t-tests with a Benjamini-Hochberg correction to contain the false discovery rate at 5% using the limma R package[24]. For multi-omic analyses, p-values for genes and proteins with matching symbols were combined with the Fisher combined probability test using the MultiGSEA R package[25,26]. Pathway enrichment was performed comparing genotypes both within each sex and with sex as an additive covariate. All expression scores for genes, proteins, and pathways are relative changes in E3KOs compared to WTs.

#### SUPPLEMENTARY RESULTS

##### Effect of Zhx2 liver overexpression (OE) on OXY CPP behaviors

We achieved a high degree of Zhx2 OE (100-fold, qPCR) in Js of both sexes via systemic AAV administration containing a cloned coding sequence of the mouse *Zhx2* gene and a thyroxine binding globulin (**TBG**) promoter to drive robust viral expression in the liver (**Fig. 1A,** plasmid map in **Fig. S6A**). We also saw no changes in brain Zhx2 expression in our first cohort of liver OE females (**Fig. 1B**), illustrating that the virus was highly selective in driving liver vs. brain OE. Liver Zhx2 OE in J females decreased locomotion across conditioning days, regardless of treatment day (**Fig. S9A**), with distance and distance on the right side being unchanged across testing days (**Fig. S9B-C**). Importantly, unlike wild-type J littermate females from the MVKO study, here, J females showed significant state-dependent CPP on Day 9 (**Fig. S9D**), thus providing an OXY phenotype with which to assess the effect of Zhx2 overexpression. Opposite to our prediction, Zhx2 liver OE increased overall time on the OXY side on all testing days but especially during the test for state-dependent OXY-CPP on Day 9. However, statistically the interaction with Day was not significant, indicating a non-specific increase in preference for the right side (**Fig. S9D**). Liver Zhx2 OE in J males also non-selectively decreased locomotion across conditioning days (**Fig. S9E**) without altering any phenotypes on testing days (**Fig. S9F-H**). Overall, liver Zhx2 OE influences locomotion irrespective of treatment day similarly in both sexes of J mice.

**SUPPLEMENTAL FIGURE LEGENDS**

**Supplemental Figure 1. Sample sizes for main figures.**

**Supplemental Figure 2. Sample sizes for supplemental figures**

**Supplemental Figure 3. Genotyping of the Zhx2 CRISPR/Cas9 lines. (A):** Zhx2 MVKO genotyping. Forward and reverse primer sequences are provided for the WT band (smaller; detects MERV presence) vs. the MVKO band (larger; detects MERV absence). Presence of both bands indicates a heterozygous MVKO. **(B):** Zhx2 E3KO genotyping. Forward and reverse primer sequences provided for WT band (detecting Exon3 presence) vs. E3KO band (detecting Exon3 absence). Presence of both bands indicates a heterozygous E3KO.

**Supplemental Figure 4. BALB/cJ *Zhx2*-MERV deletion and OXY metabolite quantification. (A):** Schematic of *Zhx2* MERV deletion in BALB/cJ mice. The deleted region began directly at the MERV insertion site and terminated directly at the end of the insertion. **(B):** Zhx2 protein quantification of all CRISPR offspring genotypes and BALB/cByJs. There was a main effect of Genotype ($F_{3,10} = 9.94$, **$p < 0.01$) and multiple comparisons indicated significant increases between MVKOs vs WTs (*$p < 0.05$), Bys vs. WTs (*$p < 0.01$), MVKOs vs. HETs (*$p < 0.05$), and Bys vs. HETs (*$p < 0.05$). **(C):** Brain OXY concentrations in MVKO females vs. WT females ($t_{19} < 1$). **(D):** Brain NOR concentrations in MVKO females vs. WT females ($t_{19} < 1$). **(E):** Brain OMOR concentrations in MVKO females vs. WT females ($t_{19} < 1$). **(F):** Brain OXY concentrations in MVKO males vs. WT males ($t_{21} < 1$). **(G):** Brain NOR concentrations in MVKO males vs. WT males ($t_{13.07} = 1.18$, $p = 0.258$). **(H):** Brain OMOR concentrations in MVKO males vs. WT males ($t_{21} < 1$).

**Supplemental Figure 5. OXY locomotion and state-dependent OXY-CPP in *Zhx2* MVKO mice. (A):** Total Distance traveled across conditioning days (Days 2-5) in females. There was a significant effect of Day ($F_{3,159} = 20.59$, ****$p < 0.0001$). However, there was no significant effects of Genotype ($F_{1,53} = 0.586$, $p = 0.447$) or Treatment ($F_{1,53} = 3.043$, $p = 0.087$), and no significant interactions involving Genotype x Treatment ($F_{1,53} = 0.135$, $p = 0.714$), Genotype x Day ($F_{3,159} = 0.362$, $p = 0.781$), Treatment x Day ($F_{3,159} = 2,911$, $p = 0.053$), or Genotype x Treatment x Day ($F_{3,159} = 0.492$, $p = 0.688$). **(B):** Total distance traveled across testing days (Days 1, 8, 9) in females. There was a main effect of Treatment ($F_{1,50} = 22.31$, ****$p < 0.0001$), Day ($F_{2,100} = 12.45$, $p < 0.0001$****) and a Treatment x Day interaction ($F_{2,100} = 43.01$, ****$p < 0.0001$). However, there was no significant effect of Genotype ($F_{1,50} = 0.064$, $p = 0.801$) and no significant interactions involving Genotype x Treatment ($F_{1,50} = 0.043$, $p = 0.836$), Genotype x Day ($F_{2,100} = 0.310$, $p = 0.681$), or Genotype x Treatment x Day ($F_{2,100} = 0.927$, $p = 0.380$). **(C):** Total distance on right traveled across testing days in females. There was a significant effect of Treatment ($F_{1,50} = 23.35$, ****$p < 0.0001$), Day ($F_{2,100} = 4.97$, *$p < 0.05$), and a Treatment x Day interaction ($F_{2,100} = 19.41$, ****$p < 0.0001$). However, there was no significant effect of Genotype ($F_{1,50} = 0.002$, $p = 0.967$) and no significant interactions involving Genotype x Treatment ($F_{1,50} = 0.003$, $p = 0.956$), Genotype x Day ($F_{2,100} = 0.746$, $p = 0.423$), and Genotype x Treatment x Day ($F_{2,100} = 0.585$, $p = 0.559$). **(D)**.

Total time on right across testing days in females. There was no significant main effect of Genotype ($F_{1,50}$ = 0.591, p = 0.446), Treatment ($F_{1,50}$ = 0.943, p = 0.336, or Day ($F_{2,100}$ = 2.481, p = 0.102). However, there were no significant interactions involving Genotype x Treatment ($F_{1,50}$ = 0.425, p = 0.518), Genotype x Day ($F_{2,100}$ = 1.263, p = 0.282), Treatment x Day ($F_{2,100}$ = 0.082, p = 0.879), or Genotype x Treatment x Day ($F_{2,100}$ = 0.513, p = 0.557). **(E):** Total distance traveled across conditioning days in males. There was a significant effect of Treatment ($F_{1,56}$ = 6.217, *p < 0.05), Day ($F_{3,168}$ = 10.81, ****p < 0.0001), and a Treatment x Day interaction ($F_{3,168}$ = 9.91, ***p < 0.001). However, there was no significant effect of Genotype ($F_{1,56}$ = 0.676, p = 0.414) and no significant interactions involving Genotype x Treatment ($F_{1,56}$ = 0.024, p = 0.879), Genotype x Day ($F_{3,168}$ = 2.140, p = 0.122), or Genotype x Treatment x Day ($F_{3,168}$ = 0.674, p = 0.513). **(F):** Total distance traveled across testing days in males. There was a significant effect of Treatment ($F_{1,53}$ = 8.68, **p < 0.01) and significant interactions involving Genotype x Day (with a visible reduction in MVKOs on Day 9; $F_{2,106}$ = 3.98, *p < 0.05), and Treatment x Day ($F_{2,106}$ = 17.15, ****p < 0.0001). However, there was no significant effect of Genotype ($F_{1,53}$ = 1.397, p = 0.243) or Day ($F_{2,106}$ = 1.966, p = 0.155), and no significant interactions involving Genotype x Treatment ($F_{1,53}$ = 0.000216, p = 0.988), or Genotype x Treatment x Day ($F_{2,106}$ = 0.723, p = 0.459). **(G):** Total distance traveled on right across testing days. There was a significant effect of Treatment ($F_{1,53}$ = 8.95, **p < 0.01) and a Treatment x Day interaction ($F_{2,106}$ = 9.37, ***p < 0.001). However, there was no significant effect of Genotype ($F_{1,53}$ = 1.947, p = 0.169) and no significant interactions involving Genotype x Treatment ($F_{1,53}$ = 0.089, p = 0.767), Genotype x Day ($F_{2,106}$ = 2.922, p = 0.075), or Genotype x Treatment x Day ($F_{2,106}$ = 0.847, p = 0.432). **(H)**: Total time on right on testing days in males. There was a significant effect of Day ($F_{2,106}$ = 6.78, **p < 0.01). However, there was no significant effect of Genotype ($F_{1,53}$ = 0.290, p = 0.593) or Treatment ($F_{1,53}$ = 0.091, p = 0.764), and no significant interactions involving Genotype x Treatment ($F_{1,53}$ = 0.359, p = 0.552), Genotype x Day ($F_{2,106}$ = 0.601, 0.525), Treatment x Day ($F_{2,106}$ = 1.311, p = 0.272), or Genotype x Treatment x Day ($F_{2,106}$ = 1.038, p = 0.349).

**Supplemental Figure 6. Zhx2 overexpression adenoassociated virus (AAV) plasmid map. (A):** AAV plasmid map for Zhx2 liver overexpression. **(B):** AAV plasmid map for Zhx2 brain overexpression.

**Supplemental Figure 7: Additional quantifications of Cyp2d expression following Zhx2 Liver OE. (A):** Cyp2d11 liver RNA expression in J females with liver OE vs. control J females ($t_{59}$ < 1). **(B):** Cyp2d10 liver RNA expression in J females with liver OE vs. control J females ($t_{27}$ < 1). **(C):** Cyp2d11 liver RNA expression in J males with liver OE vs. control J males ($t_{42}$< 1). **(D):** Cyp2d10 liver RNA expression in J males with liver OE vs. control J males ($t_{42}$ = 1.63, p = 0.110).

**Supplemental Figure 8: OXY and NOR quantification following Zhx2 Liver OE. (A):** Brain OXY concentrations in J females with liver OE vs control J females ($t_{46}$ < 1). **(B):** Brain NOR concentrations in J females with liver OE vs control J females ($t_{46}$ < 1). **(C):** Brain OXY concentrations in J males with liver OE vs. control J males ($t_{44}$ = 2.12, *p < 0.05). **(D):** Brain NOR concentrations in J males with liver OE vs. control J males ($t_{44}$ < 1).

**Supplemental Figure 9. OXY locomotion and state-dependent OXY-CPP following Zhx2 Liver OE. (A):** Total distance traveled across conditioning days in females. There was a significant effect of AAV ($F_{1,54}$ = 6.33, *p < 0.05) and Day ($F_{3,162}$ = 64.21, ****p < 0.0001), but no significant AAV x Day interaction ($F_{3,162}$ = 2.244, p = 0.109). **(B):** Total distance traveled across testing days in females. There was a significant effect of Day ($F_{2,94}$ = 59.28, ****p < 0.0001), but no significant effect of AAV ($F_{1,47}$ = 0.414, p = 0.523) or a significant AAV x Day interaction ($F_{2,94}$ = 0.178, p = 0.752). **(C):** Total distance traveled on right across testing days in females. There was a significant effect of Day ($F_{2,94}$ = 50.52, ****p < 0.0001), but no significant effect of AAV ($F_{1,47}$ = 3.583, p = 0.065) or significant AAV x Day interaction ($F_{2,94}$ = 2.905, p = 0.091). **(D)**. Total time on right traveled across testing days in females. There was a significant effect of AAV ($F_{1,47}$ = 7.65, **p < 0.01) and Day ($F_{2,94}$ = 15.79, ****p < 0.0001) but no significant AAV x Day interaction ($F_{2,94}$ = 1.913, p = 0.170). **(E):** Total distance traveled across conditioning days in males. There was a significant effect of AAV ($F_{1,46}$ = 4.14, *p < 0.05) and Day ($F_{3,138}$ = 18.41, ****p < 0.0001), but no significant AAV x Day interaction ($F_{3,138}$ = 1.325, p = 0.271). **(F):** Total distance traveled across testing days in males. There was a significant effect of Day ($F_{2,84}$ = 20.99, ****p < 0.0001), but no significant effect of AAV ($F_{1,42}$ = 2.151, p = 0.150) or a significant AAV x Day interaction ($F_{2,84}$ = 0.333, p = 0.667). **(G):** Total distance right traveled across testing days. There was a significant effect of Day ($F_{2,84}$ = 38.49, ****p < 0.0001), but no significant effect of AAV ($F_{1,42}$ = 0.922, p = 0.342) or a significant AAV x Day interaction ($F_{2,84}$ = 0.017, p = 0.949). **(H)**: Total time on right traveled on testing days in males. There was a significant effect of Day ($F_{2,84}$ = 12.75, ***p < 0.001), but no significant effect of AAV ($F_{1,42}$ = 2.889, p = 0.097) or a significant AAV x Day interaction ($F_{2,84}$ = 0.425, p = 0.591).

**Supplemental Figure 10. Image locations for brain Zhx2 OE.**
Order from rostral to caudal: Lateral Septum (Rostral), Bed Nucleus of the Stria Terminalis, Lateral Septum (Caudal), CA3 (Rostral), CA3 (Caudal).

**Supplemental Figure 11. Differential expression analysis in the brain of *Zhx2* KO mice. (A):** Differentially expressed genes in E3KO females vs. WT females using RNA-Seq. **(B):** Differentially expressed proteins in E3KO females vs. WT females using mass-spectrometry. **(C):** Differentially expressed genes in E3KO females vs. WT females using RNA-Seq. **(D):** Differentially expressed proteins in E3KO males vs. WT males using mass-spectrometry.

**Supplemental Figure 12. Sex-specific differential gene/protein expression in the brain and pathway enrichment in Zhx2 E3KO mice. (A):** Top differentially expressed proteins in one sex (p.adj < 0.05) that were not differentially expressed in the other (p.unadj > 0.05). **(B):** Top differentially expressed genes/proteins from combined -omics in one sex (p.adj < 0.05) that were not differentially expressed in the other (p.unadj > 0.05). **(C):** Top enriched pathways from genes differentially expressed in E3KO females (p.unadj < 0.05) but not males (p.unadj > 0.05). **(D):** Top enriched pathways from genes differentially expressed in E3KO males (p.unadj < 0.05) but not females (p.unadj > 0.05).

**Supplemental Figure 13. Sex- and -omics collapsed leading edge gene/protein interactions from top pathways in Zhx2 KO mice.** STRING-db plot graphing interactions between top 10 unique leading-edge genes/proteins from top 8 pathways. Colors indicate clusters based on a Markov Clustering Algorithm, with line thickness indicating strength of interaction based on existing data resources.

**Supplemental Table 1. Antibodies.**

**Supplemental Table 2. qPCR primers.**

**Supplemental Table 3. RNA-seq gene list (Genotype - Sex covariate).**

**Supplemental Table 4. RNA-seq gene list (Genotype - Males).**

**Supplemental Table 5. RNA-seq gene list (Genotype - Females).**

**Supplemental Table 6. Mass-spectrometry protein list (Genotype - Sex covariate).**

**Supplemental Table 7. Mass-spectrometry protein list (Genotype - Males).**

**Supplemental Table 8. Mass-spectrometry protein list (Genotype - Females).**

**Supplemental Table 9. Multi-omics gene/protein list (Genotype - Sex covariate).**

**Supplemental Table 10. Multi-omics gene/protein list (Genotype - Males).**

**Supplemental Table 11. Multi-omics gene/protein list (Genotype - Females).**

**Supplemental Table 12. RNA-seq pathway enrichment list (Genotype - Sex covariate).**

**Supplemental Table 13. RNA-seq pathway enrichment list (Genotype - Males).**

**Supplemental Table 14. RNA-seq pathway enrichment list (Genotype - Females).**

**Supplemental Table 15. Mass-spectrometry pathway enrichment list (Genotype - Sex covariate).**

**Supplemental Table 16. Mass-spectrometry pathway enrichment list (Genotype - Males).**

**Supplemental Table 17. Mass-spectrometry pathway enrichment list (Genotype - Females).**

**Supplemental Table 18. Multi-omics pathway enrichment list (Genotype - Sex covariate).**

**Supplemental Table 19. Multi-omics pathway enrichment list (Genotype - Males).**

**Supplemental Table 20. Multi-omics pathway enrichment list (Genotype - Females).**

**Supplemental Table 21. MCL clustering of top leading-edge gene/proteins.**

**SUPPLEMENTAL FIGURE 1**

| Figure 1 | | |
|---|---|---|
| | CTR | Liver OE |
| 1A | 53 | 52 |
| 1B | 8 | 8 |
| 1C | 30 | 31 |
| 1D | 24 | 24 |
| 1E | 23 | 21 |
| 1F | 23 | 23 |

| Figure 3 | | | | |
|---|---|---|---|---|
| | WT | HET | E3KO | J |
| 3B | 4 | 4 | 4 | 3 |
| 3C-E | 13 | | 13 | |
| 3F-H | 7 | | 7 | |

| Figure 2 | | |
|---|---|---|
| | CTR | Brain OE |
| 2A | 16 | 16 |
| 2B-E | 16 | 15 |
| 2F | 16 | 16 |
| 2G-J | 16 | 16 |

| Figure 4 | | | | |
|---|---|---|---|---|
| | WT (SAL) | WT (OXY) | E3KO (SAL) | E3KO (OXY) |
| 4A | 14 | 15 | 13 | 14 |
| 4B-E | 15 | 15 | 13 | 14 |
| 4F | 7 | 10 | 8 | 11 |
| 4G-J | 7 | 9 | 9 | 10 |

| Multi-Omics | | | | |
|---|---|---|---|---|
| | WT - M | WT - F | E3KO - M | E3KO - F |
| RNA | 4 | 4 | 4 | 4 |
| Protein | 4 | 4 | 4 | 4 |

#### SUPPLEMENTAL FIGURE 2

| Figure S1 | | | | |
|---|---|---|---|---|
| | WT | HET | MVKO | By |
| S1B | 3 | 3 | 4 | 4 |
| S1C-E | 16 | | 5 | |
| S1F-H | 12 | | 11 | |

| Figure S2 | | | | |
|---|---|---|---|---|
| | WT (SAL) | WT (OXY) | MVKO (SAL) | MVKO (OXY) |
| S2A | 14 | 20 | 11 | 12 |
| S2B-D | 15 | 18 | 10 | 11 |
| S2E | 15 | 17 | 13 | 15 |
| S2F-H | 15 | 15 | 13 | 14 |

| Figure S5 | | |
|---|---|---|
| | CTR | Liver OE |
| S5A | 30 | 31 |
| S5B | 16 | 13 |
| S5C | 23 | 21 |
| S5D | 23 | 21 |

| Figure S6 | | |
|---|---|---|
| | CTR | Liver OE |
| S6A-B | 24 | 24 |
| S6C-D | 23 | 23 |

| Figure S7 | | |
|---|---|---|
| | CTR | Liver OE |
| S7A | 28 | 28 |
| S7B-D | 25 | 24 |
| S7E | 23 | 25 |
| S7F-H | 23 | 21 |

#### SUPPLEMENTAL FIGURE 3

### MVKO

Primers(WT band):

*Forward:* ACTGTCTCAGCTCATTCCCTGCAA

*Reverse:* CCAGTAGAAAGGTGCACGGGT

Primers(MERV-KO band):

*Forward:* ACTGTCTCAGCTCATTCCCTGCAA

*Reverse:* AATGCTTCACATGGCACACAGCAG

Expected Band Sizes:
WT: 283 bp
MERV-KO: 342 bp

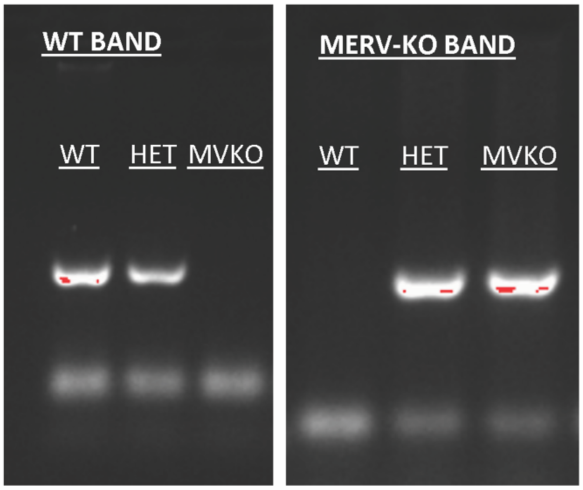


# E3KO

Primers(WT band):

*Forward:* TCAAGCGCAATAATCAGACG

*Reverse:* CTTTCTTGGCATCTGCCTTC

Primers(Exon3-KO band):

*Forward:* TTTGTGTCTCTGAGCATGGAG

*Reverse:* CCTTTCCTTCTTGGGTTTGT

Expected Band Sizes:
WT: 165 bp
Exon3-KO: 294 bp

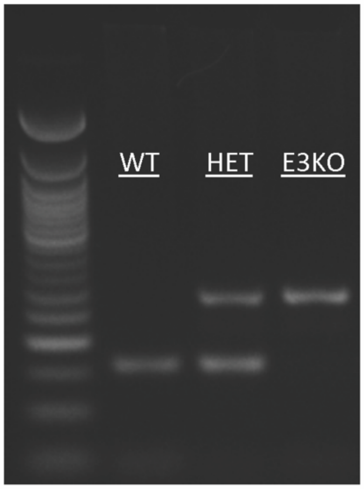

**SUPPLEMENTAL FIGURE 4**

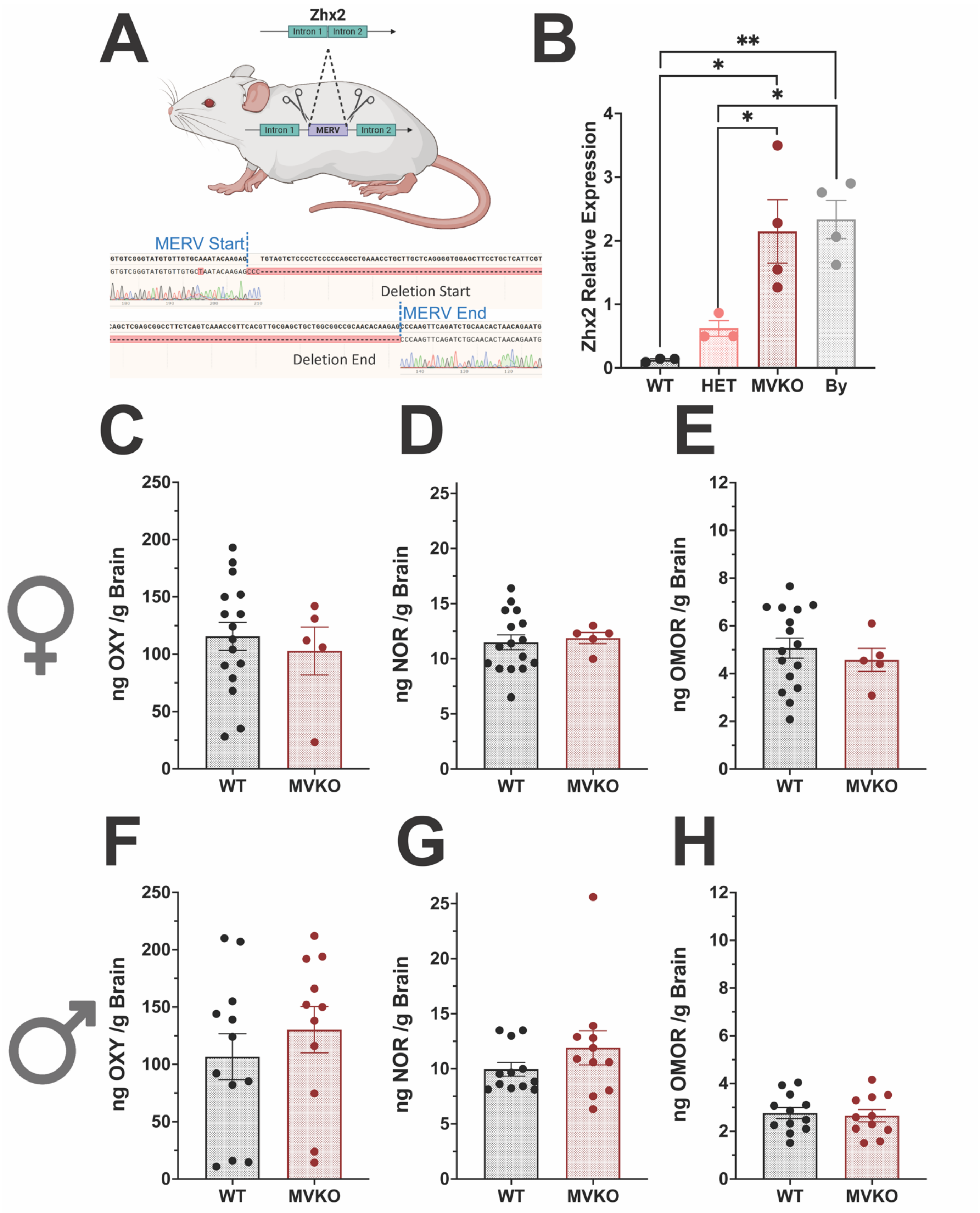

**SUPPLEMENTAL FIGURE 5**

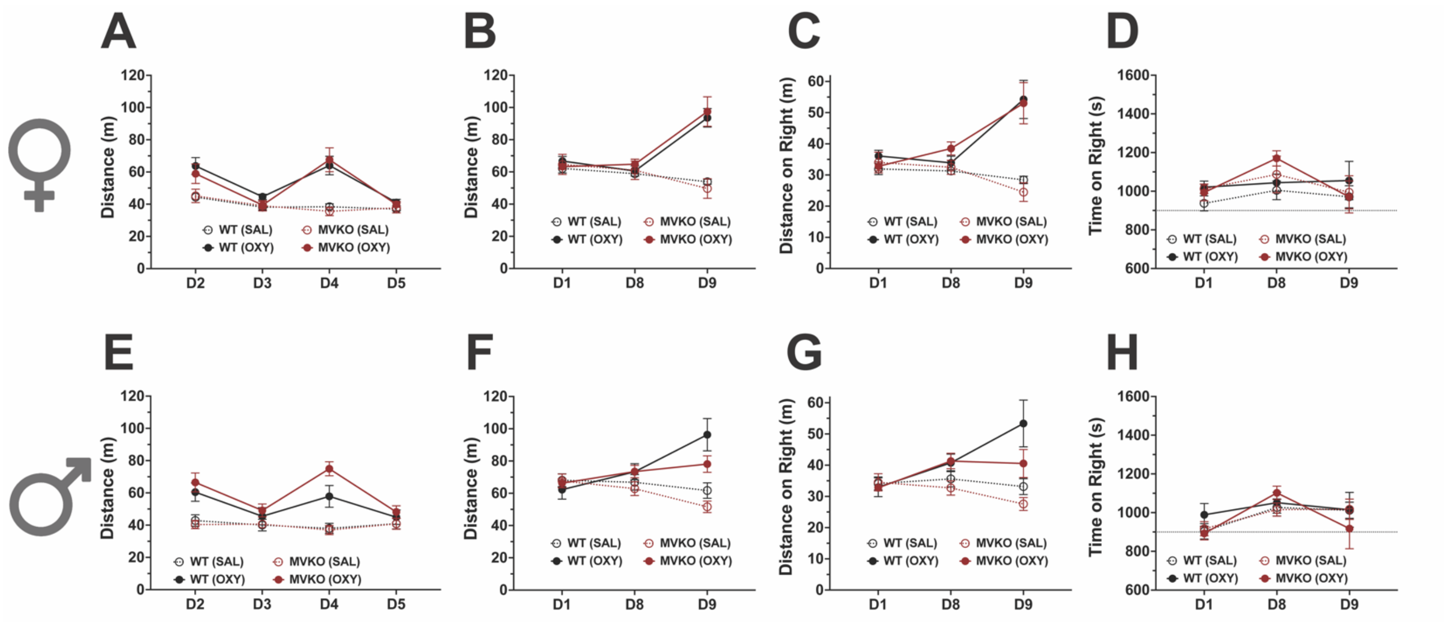

#### SUPPLEMENTAL FIGURE 6

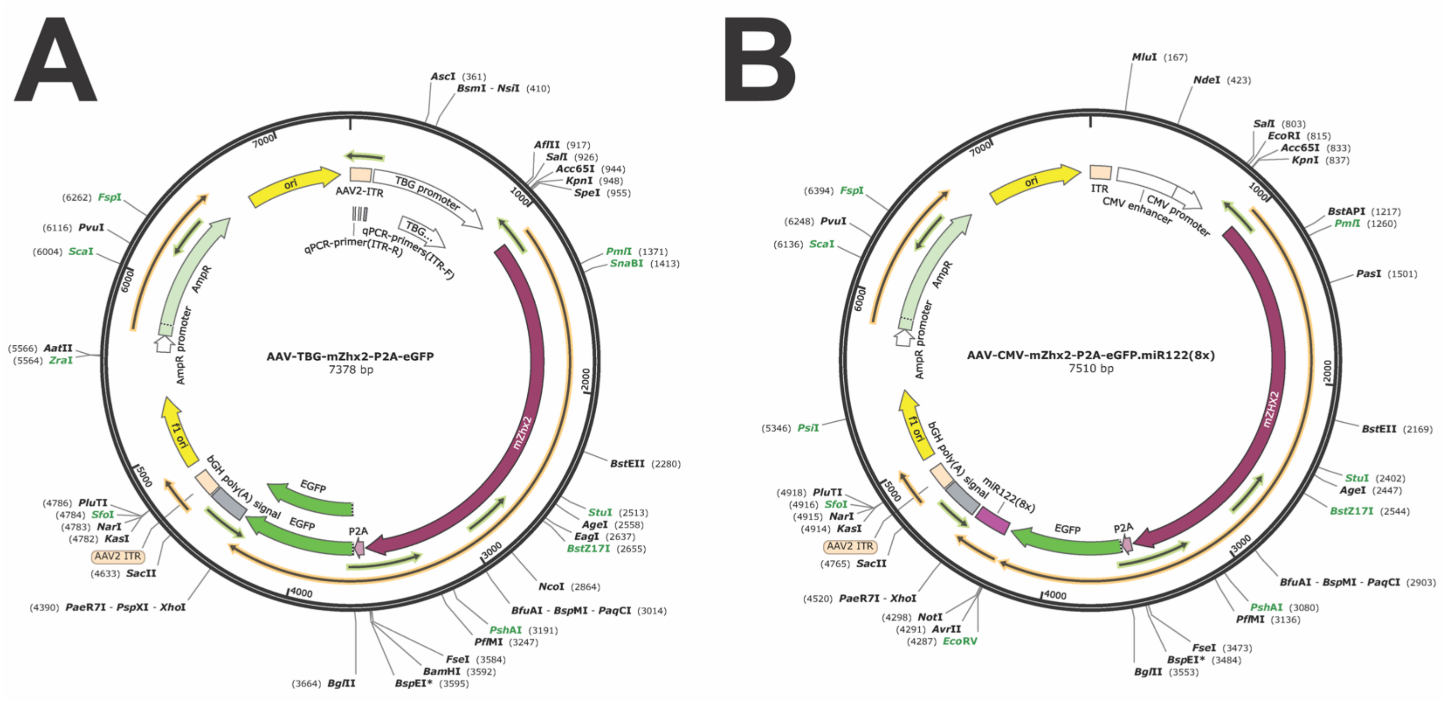

**SUPPLEMENTAL FIGURE 7**

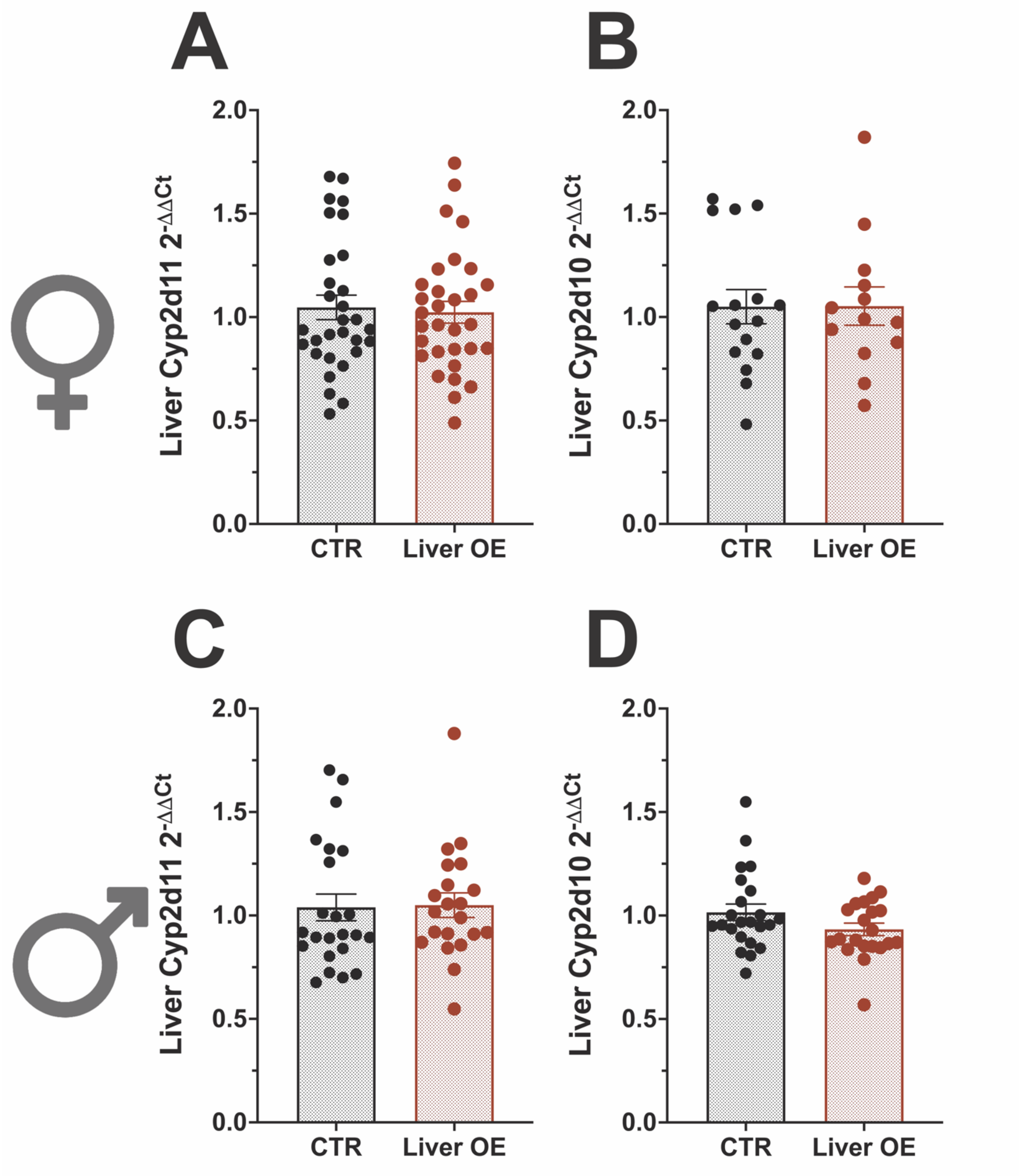

**SUPPLEMENTAL FIGURE 8**

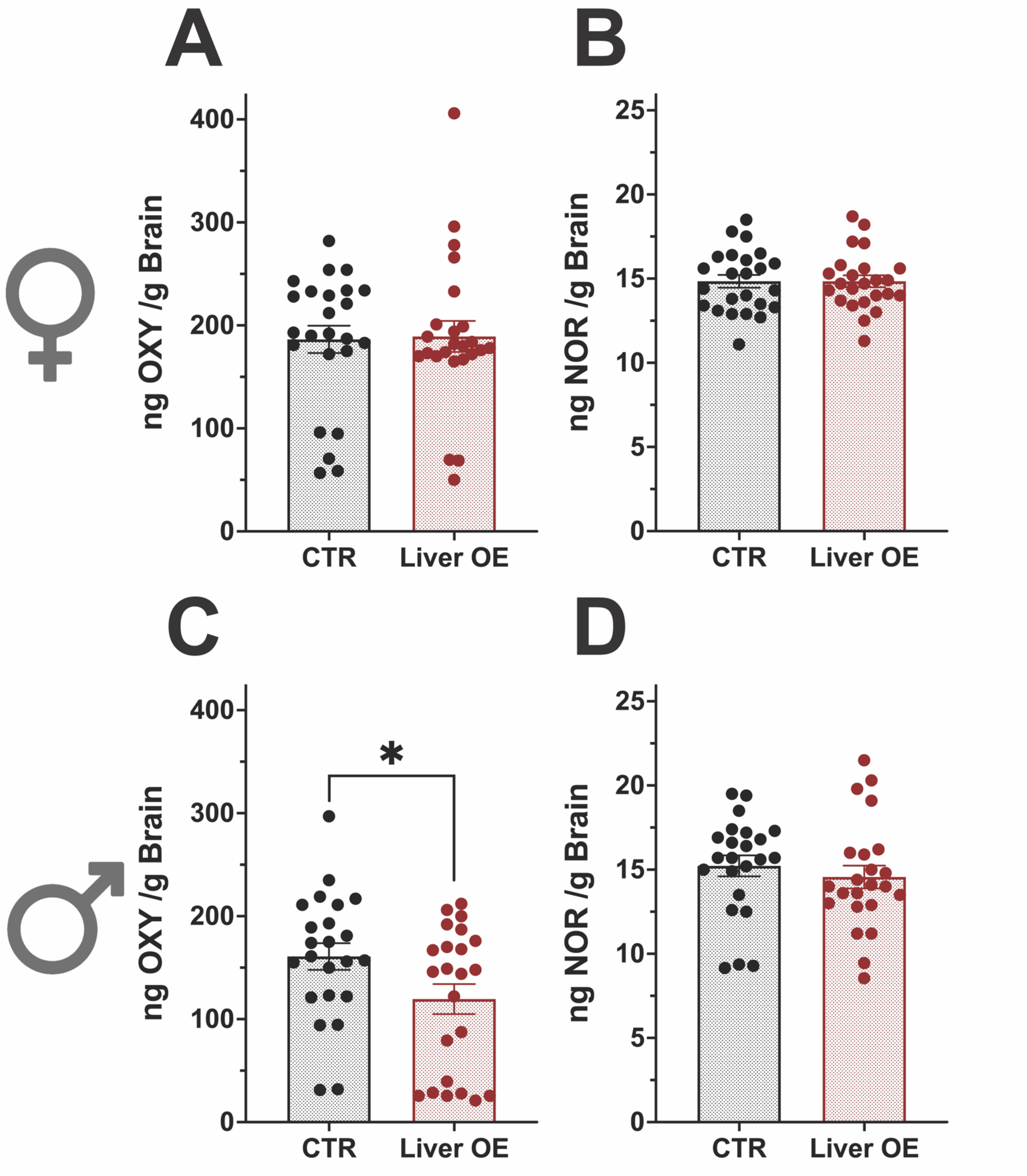

**SUPPLEMENTAL FIGURE 9**

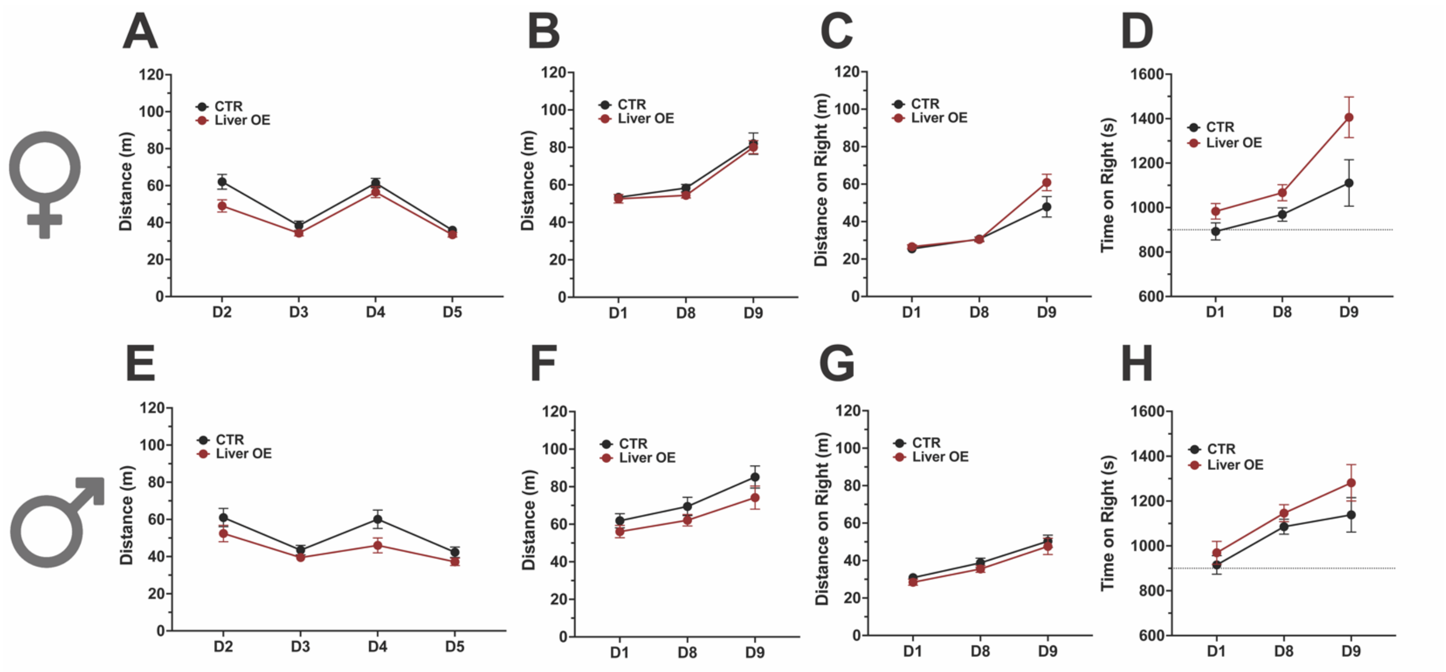

SUPPLEMENTAL FIGURE 10

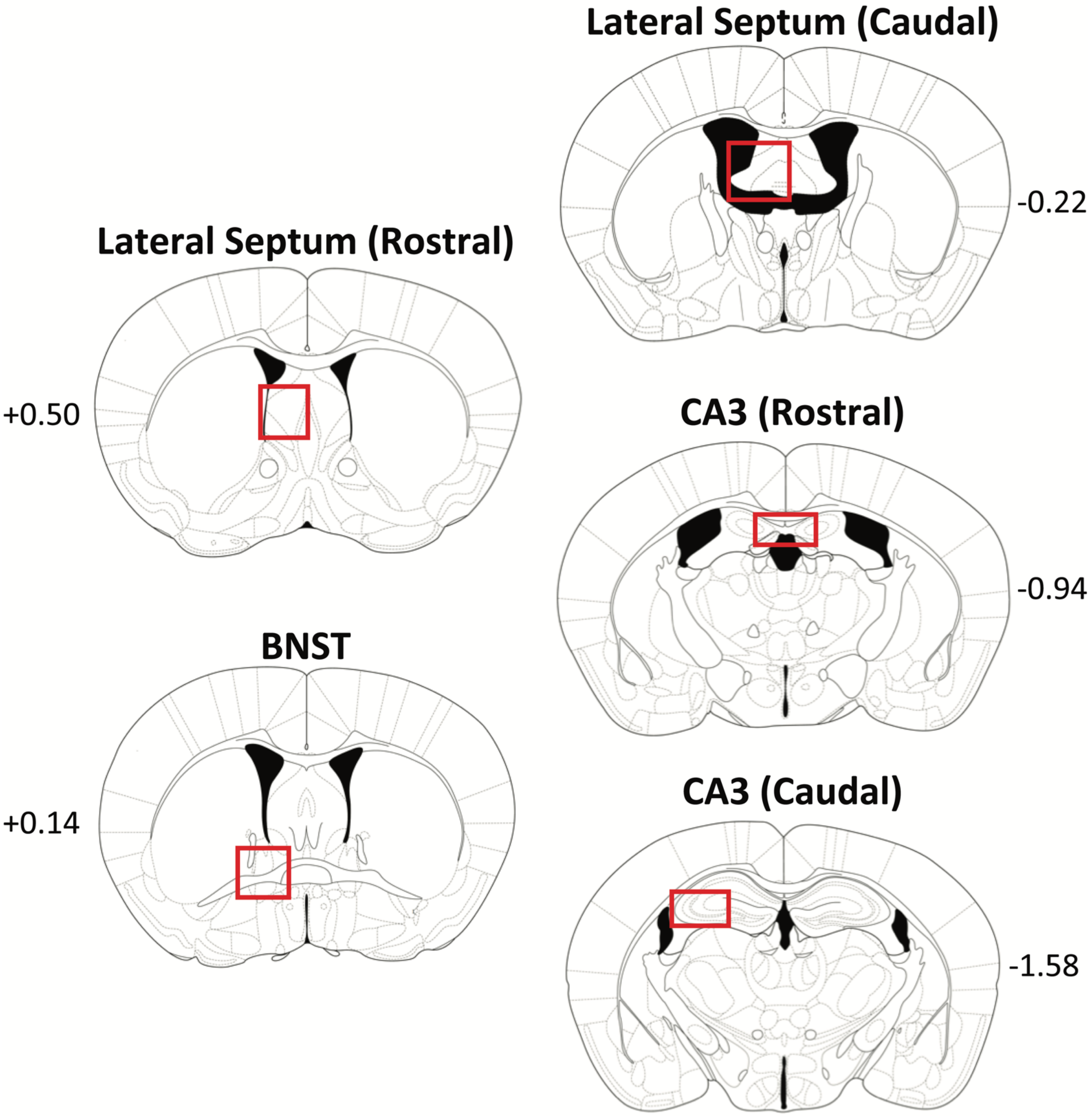

**SUPPLEMENTAL FIGURE 11**

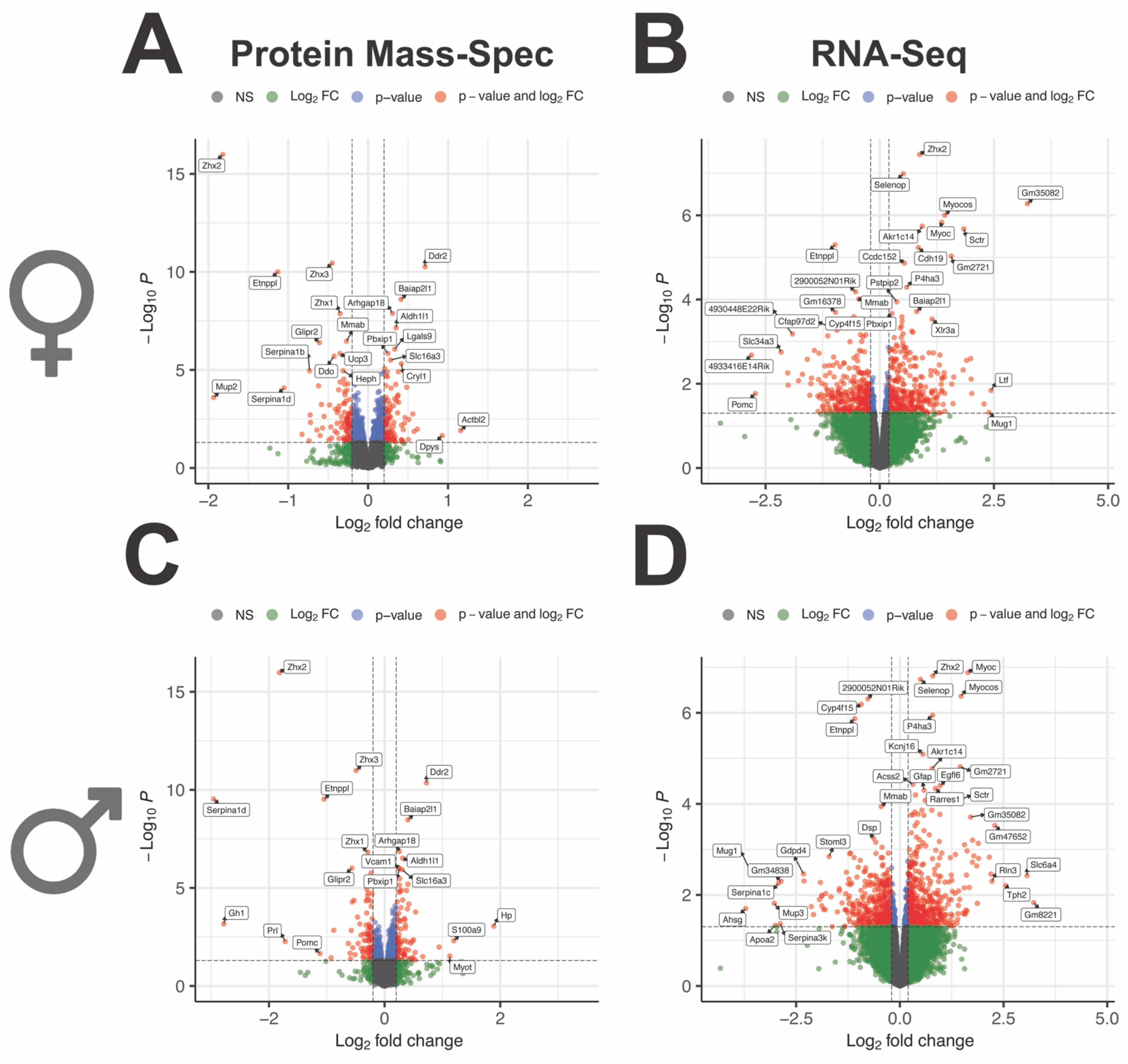

**SUPPLEMENTAL FIGURE 12**

**A**

**PROTEIN**

| Gene Symbol | Sex | P.value | P.adj |
|---|---|---|---|
| Dennd3 | Male | 8.26E-05 | 0.02364625 |
| Dhtkd1 | Male | 0.000113721 | 0.029980427 |
| Elovl7 | Male | 0.000168874 | 0.041262855 |
| Trim23 | Male | 0.00017749 | 0.04233569 |
| Serpina1b | Female | 1.11E-05 | 0.005835429 |
| Grpel2 | Female | 1.28E-05 | 0.005980517 |
| Dapk2 | Female | 2.21E-05 | 0.008867538 |
| Mid1 | Female | 5.14E-05 | 0.017760473 |
| Sez6l | Female | 5.87E-05 | 0.018366426 |
| Apoa4 | Female | 0.000103118 | 0.027185245 |
| Rbm8a | Female | 0.000203756 | 0.044374613 |
| Slco1c1 | Female | 0.00021523 | 0.04587597 |

**B**

**Combined**

| Gene Symbol | Sex | P.value | P.adj |
|---|---|---|---|
| Pon2 | Male | 3.85E-05 | 0.010182386 |
| Dhtkd1 | Male | 0.000230616 | 0.037464987 |
| Cyp4f14 | Male | 0.00028804 | 0.044454156 |
| Dapk2 | Female | 8.25E-06 | 0.003321194 |
| Mid1 | Female | 2.21E-05 | 0.006592429 |
| Scrg1 | Female | 6.23E-05 | 0.014802536 |
| Grpel2 | Female | 7.54E-05 | 0.01702878 |
| Serpina1b | Female | 0.000105582 | 0.022737047 |

**C**

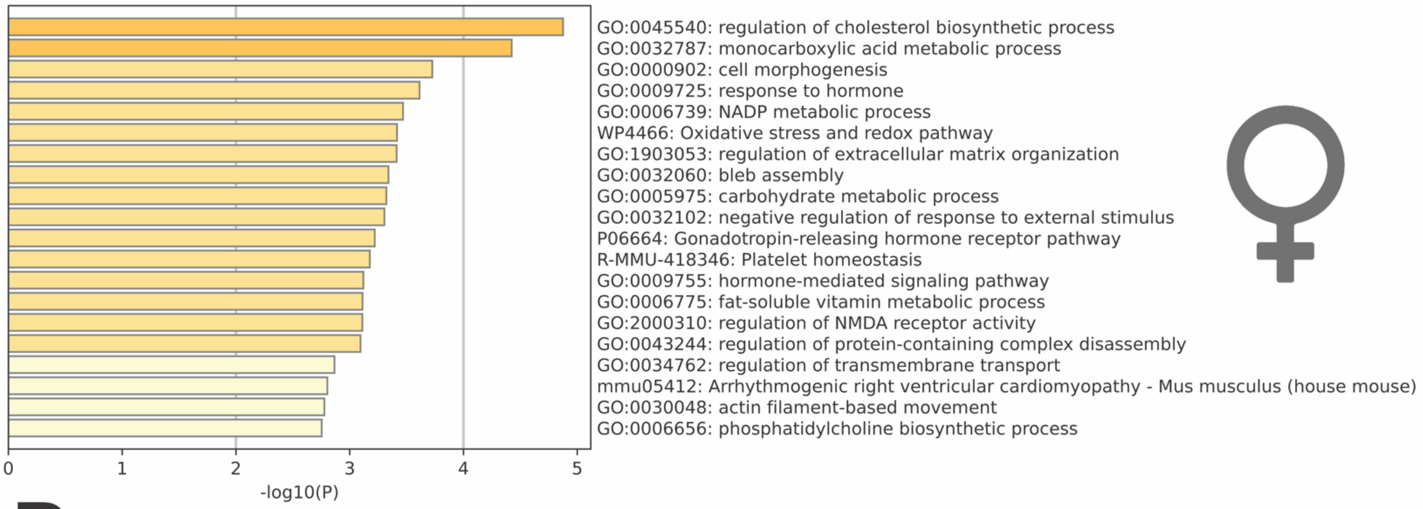


**D**

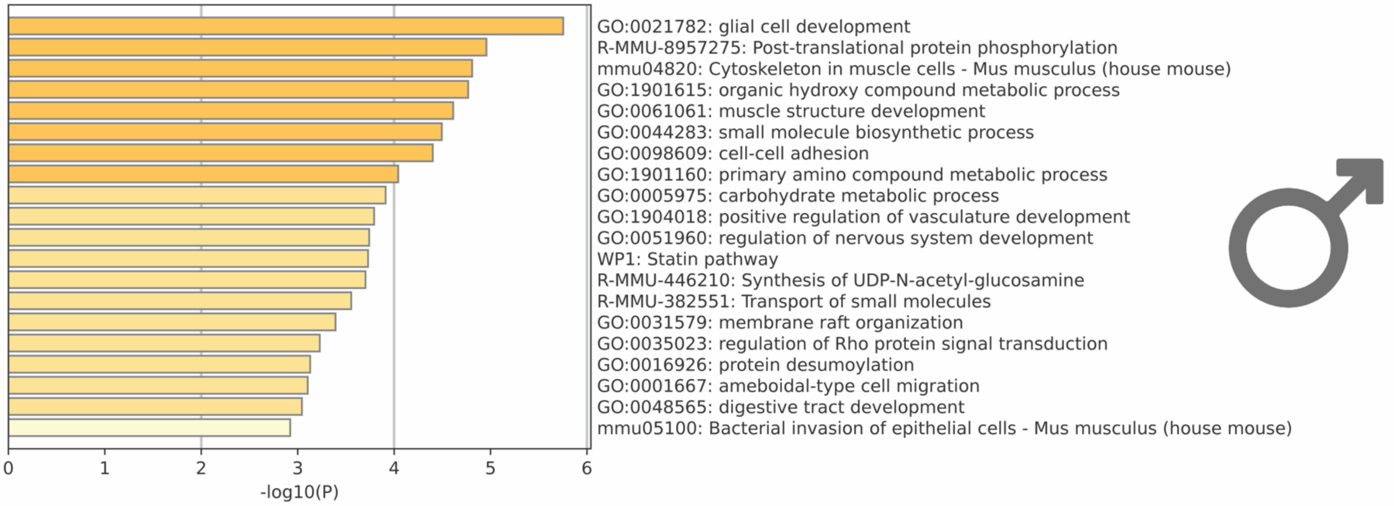

**SUPPLEMENTAL FIGURE 13**

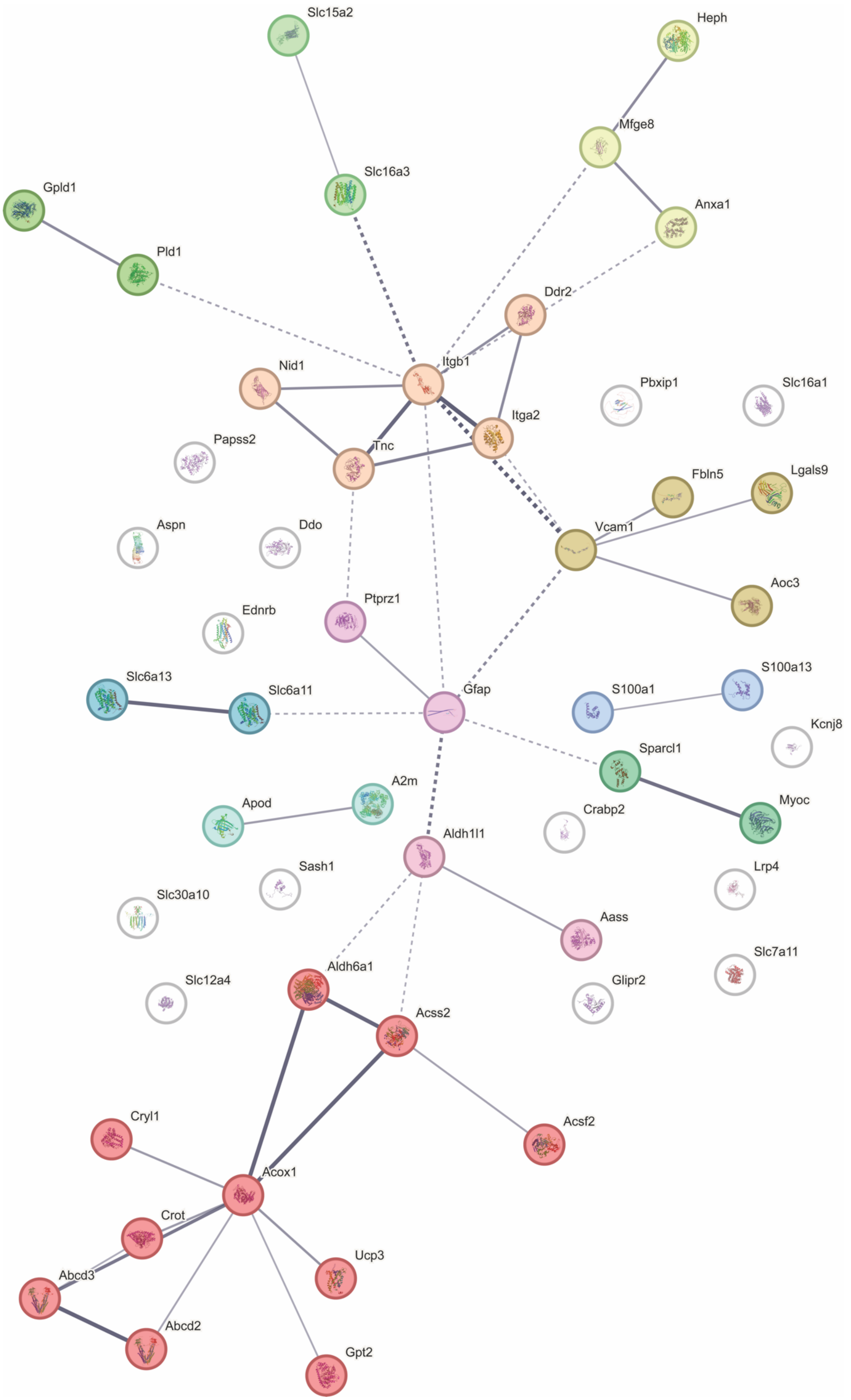

#### SUPPLEMENTAL TABLE 1

| Antibody | Type | Use | Concentration | Product Number | Company |
|---|---|---|---|---|---|
| ZHX2 antibody [C1C3] (Rabbit Polyclonal) | Primary | Western Blotting | 1:4000 | GTX112232 | GeneTex |
| Peroxidase-conjugated AffiniPure Donkey Anti-Rabbit | Secondary | Western Blotting | 1:10K | 711-035-152 | Jackson ImmunoResearch |
| ZHX2 antibody [C1C3] (Rabbit Polyclonal) | Primary | IHC | 1:750 | GTX112232 | GeneTex |
| Donkey anti-Rabbit IgG Alexa Fluor™ 568 | Secondary | IHC | 1:750 | A10042 | ThermoFisher |

**SUPPLEMENTAL TABLE 2**

| Primer Name | Sequence | Source |
|---|---|---|
| Zhx2_Forward | GTTCTTTGCCCAGTCCTTC | Song et al. 2018 |
| Zhx2_Reverse | CGCCTCGCTTCCATTCTT | Song et al. 2018 |
| Cyp2d22_Forward | GGGCCTTTGTTACCATGTTGG | Blume et al. 2000 |
| Cyp2d22_Reverse | TACTCGGCGCTGCACATCTG | Blume et al. 2000 |
| Cyp2d11_Forward | TCTCAGTGCCTGATGGACAG | Zhong et al. 2018 |
| Cyp2d11_Reverse | CACAGAGCTGGTAGGGGAAG | Zhong et al. 2018 |
| Cyp2d10_Forward | TCCACTGAATTTGCCACGC | Elraghy and Baldwin 2015 |
| Cyp2d10_Reverse | TCAGCACGGAGGACATGTTG | Elraghy and Baldwin 2015 |
| Gapdh_Forward | GCCTTCCGTGTTCCTACC | Ruan et al. 2022 |
| Gapdh_Reverse | CCTCAGTGTAGCCCAAGATG | Ruan et al. 2022 |
